# Pattern-based genome mining guides discovery of the antibiotic indanopyrrole A from a marine streptomycete

**DOI:** 10.1101/2024.10.29.620887

**Authors:** Douglas Sweeney, Alexander Bogdanov, Alexander B. Chase, Gabriel Castro-Falcón, Alma Trinidad-Javier, Samira Dahesh, Victor Nizet, Paul R. Jensen

**Author notes:** D.S. and A.B. contributed equally to this work. The manuscript was written through contributions of all authors. All authors have given approval to the final version of the manuscript.

## Abstract

Terrestrial actinomycetes in the genus *Streptomyces* have long been recognized as prolific producers of small molecule natural products, including many clinically important antibiotics and cytotoxic agents. Although *Streptomyces* can also be isolated from marine environments, their potential for natural product biosynthesis remains underexplored. The MAR4 clade of largely marine-derived *Streptomyces* has been a rich source of novel halogenated natural products of diverse structural classes. To further explore the biosynthetic potential of this group, we applied pattern-based genome mining leading to the discovery of the first halogenated pyrroloketoindane natural products, indanopyrrole A (**1**) and B (**2**), and the bioinformatic linkage of these compounds to an orphan biosynthetic gene cluster (BCG) in 20 MAR4 genomes. Indanopyrrole A displays potent broad-spectrum antibiotic activity against clinically relevant pathogens. A comparison of the putative indanopyrrole BGC with that of the related compound indanomycin provides new insights into the terminal cyclization and offloading mechanisms in pyrroloketoindane biosynthesis. Broader searches of public databases reveal the rarity of this BGC while also highlighting opportunities for discovering additional compounds in this uncommon class.

**GRAPHICAL ABSTRACT:** 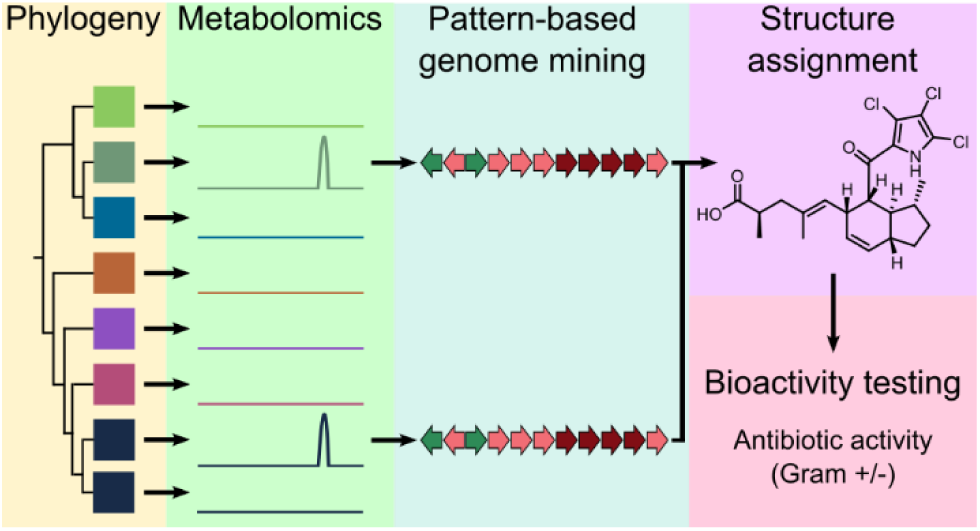

## INTRODUCTION

Twenty-eight percent of drugs approved by the FDA in 2021 contained at least one halogen atom.^1^ The presence of these electronegative atoms can enhance compound bioactivity^2^ as observed in the antibiotic vancomycin^3^ and the anticancer drug salinosporamide A.^4^ The biological activity of halogenated natural products makes them attractive targets for drug discovery, and the abundance of halogen atoms in the marine environment offers a unique opportunity for discovering new halogenated metabolites. In 2015, it was estimated that more than 5,000 halogenated natural products had been discovered, with the majority produced by marine organisms, including bacteria.^5^ Among marine bacteria, the MAR4 group of marine-derived *Streptomyces* have yielded a wide diversity of halogenated natural products^6^ including phenazines,^7^ nitropyrroles,^8,9^ and tetrahydroxynaphthalene (THN)-derived molecules.^10–14^ In addition to these diverse halogenated compounds, genome mining has revealed halogenases in orphan MAR4 biosynthetic gene clusters (BGCs), suggesting that additional halogenated metabolites await discovery.^15,16^

One approach to facilitate natural product discovery involves the pairing of genomic and metabolomic data collected from closely related strains, a process known as pattern-based genome mining or metabologenomics.^17,18^ This method correlates gene cluster families with ions detected by mass spectrometry (MS), enabling the connection of metabolites to orphan BGCs, as shown in the marine actinomycete *Salinispora*^17^ and other bacteria.^18^ The technique is also applicable to BGC subclusters, as shown for the pyrrole-containing compounds chlorizidine,^19^ armeniaspirol,^20^ and marinopyrrole.^21^ Recent advances in pattern-based genome mining include automated tools such as NPLinker^22^ and NPOmix,^23^ which use correlation-based statistics and machine learning to computationally link metabolites to their cognate BGCs. Halogenated natural products are especially well suited for pattern-based genome mining due to their distinct isotopic signatures and biosynthetic genes that can be used as ‘hooks’ to detect candidate BGCs. Linking metabolomic and biosynthetic sequence data in this way can facilitate the discovery of novel halogenated metabolites, with deeper BGC analyses facilitating structure assignments.^24^

Pyrroloketoindanes are characterized by indane and pyrrole systems bridged via a ketone. Compounds containing these ring systems are rare and often biologically active, making them logical targets for natural product discovery. Prior to this study, only six pyrroloketoindanes had been reported, none of which were halogenated (Chart 1). These compounds, all of bacterial origin, are the products of polyketide synthase (PKS) BGCs. The first pyrroloketoindane described was indanomycin, which contains a tetrahydropyran and was discovered in 1979 from *Streptomyces antibioticus* NRRL 8167^25^. Indanomycin has activity against Gram-positive bacteria^26^ and acts as an ionophore. ^27,28^ Later work has shown that indanomycin has insecticidal^29^ and antiviral activities.^30^ The indanomycin analogue cafamycin was identified from a *Streptomyces* species in 1987.^31^ Shortly thereafter, 16-deethylindanomycin was isolated from *Streptomyces setonii* A80394A and reported to be active against Gram-positive bacteria and protist parasites.^32^ A compound with the same structure as 16-deethylindanomycin was published in 1990 under the name omomycin and reported to elevate cyclic guanosine monophosphate levels in rat heart cells.^33^ The pyrroloketoindane containing compound homoindanomycin was described in a patent from 1989 without any information about activity.^34^ Stawamycin inhibits Epstein-Barr viral transcription factor BZLF1 binding to DNA and was the first indanomycin analogue described to lack a tetrahydropyran.^35^ Finally, JBIR-11 was reported in 2008 from *Streptomyces viridochromogenes* subsp. *sulfomycini* NBRC 13830. This unusual analogue of stawamycin contains a tryptophan moiety and was shown to be cytotoxic to HT-1080 fibrosarcoma cells.^36^

Here we describe the discovery of novel di- and trichlorinated pyrroloketoindane antibiotics using paired metabolomic and genomic datasets. The candidate BGC provides new insights into pyrroloketoindane biosynthesis while its distribution in bacterial genomes reveals opportunities for additional compound discovery.

## RESULTS AND DISCUSSION

### Isolation and structure elucidation

A search for novel, halogenated natural products from *Streptomyces* strains in the MAR4 clade led to the detection of a compound with an *m/z* of 458.1060 [M+H]^+^ and an isotopic pattern characteristic of a trichlorinated molecule in the culture extract of strain CNX-425 (Figure S1). A Dictionary of Natural Products database search retrieved no matches, suggesting it represented a new compound. The compound, which we have named indanopyrrole A (**1**), was isolated and structurally characterized as described below. During the isolation of **1**, we obtained and characterized a minor related compound (**2**) that shared the same chromophore and was named indanopyrrole B (Chart 1).

### Chart 1

Indanopyrroles A and B (**1**-**2**) and previously reported pyrroloketoindane natural products. Pyrroloketoindane moieties are colored green. Absolute configurations for **1-2** are based on bioinformatic prediction.

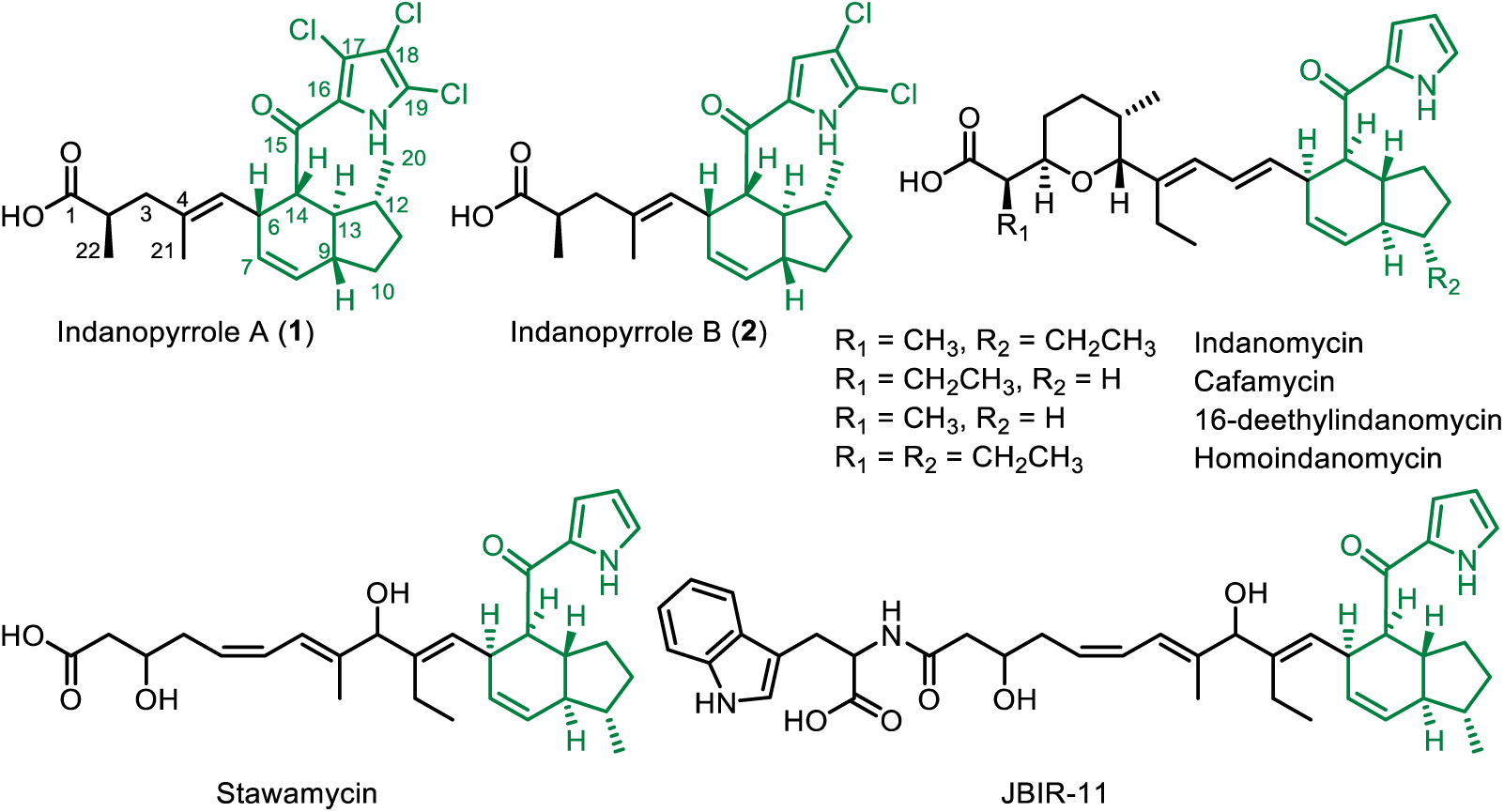

The molecular formula (MF) of **1** C_22_H_26_Cl_3_NO_3_ with nine degrees of unsaturation was determined from the accurate mass (HR-ESI-TOF-MS) measurement. NMR experiments (^1^H, ^13^C, COSY, HSQC and HMBC) in CD_3_OD led to the unambiguous elucidation of the planar structure (Table S1). The proton spectrum showed the presence of two methyl doublets (CH_3_-20 and CH_3_- 22; δH 0.99, *J* = 6.1 Hz and 0.98, *J* = 6.8 Hz) and one methyl singlet (CH_3_-21; δH 1.31) attached to an olefin. Additionally, three *sp^2^*resonances at 5.06 (H-5, d, *J* = 10.3 Hz), 5.33 (H-7, dt, *J* = 9.7, 3.4 Hz), and 5.84 ppm (H-8, dt, *J* = 9.7, 1.8 Hz) accounted for two olefins. In the ^1^H-^1^H COSY spectrum, an extensive spin system from H-5 to H-14 established the tetrahydroindane core of the molecule (Figure S2A), the position of methyl C-12, and two substituents at C-6 and C-14. The sidechain at C-6 was established using ^1^H-^13^C HMBC, which showed long range correlations between the methyl protons H_3_-21 to C-3, C-4, and C-5 and from H_3_-22 to the carboxylic acid carbonyl C-1 (δC 182.7 ppm), C-2, and C-3. The substructure composed of the tetrahydroindane core and the C-6 substituent accounted for 17 carbons, 25 protons, two oxygens, and five degrees of unsaturation. The remaining C_5_HCl_3_NO was suggested to be a trichlorinated pyrrole- 2-carbonyl substituent on C-14 based on a fragment of 195.9127 *m/z* in the MS/MS spectrum of **1** (Figure S3).^32,35–37^ The UV maximum absorbance at 295 nm provided further support for the pyrrole moiety. HMBC correlations from H-14 to the carbonyl C-15 and a sp^2^ hybridized carbon C-16 (δC 128.0 ppm) established the connection of the trichloropyrrole moiety to the indane core via a ketone bridge. Analysis of the ^13^C NMR spectrum revealed the chemical shifts for the remaining chlorinated carbons (C-17, C-18 and C-19) that were not observed by indirect detection experiments (HSQC and HMBC) and secured the planar structure of indanopyrrole A (**1**, Chart 1). The configuration of the C-4, C-5 olefin was assigned as *trans* (*E*) based on the shielded ^13^C resonance of the methyl substituent CH_3_-21 (δC 15.7 ppm) and on a clear NOE correlation between H-5 and H-3a. The relative configuration of the five stereocenters on the indane was established using NOESY experiment (NOE correlations observed between H-6, H-9 and H-14, as well as from H-13 to H_3_-20) and is in accordance with previously reported pyrroloketoindanes (Chart 1). The absolute configurations at C-2 and C-12 in the linear biosynthetic precursor were predicted bioinformatically as 2*R* and 12*S* (12*R* after indane formation) based on an analysis of the candidate biosynthetic gene cluster (see below). This, in turn, led to the absolute configuration of all stereocenters in **1** (2*R*, 6*R*, 9*R*, 12*R*, 13*S*, 14*R* Figure S3B-C) and differences in the indane core compared to other pyrroloketoindanes (Chart 1). Raw NMR data files can be accessed from the Natural Product Magnetic Resonance Database Project (<u>NP-MRD</u> https://np-mrd.org/) under identifiers NP0341895 (**1**) and NP0341896 (**2**). HR-ESI-TOFMS/MS analyses of **2** revealed the molecular formula C_22_H_27_Cl_2_NO_3_ and a prominent dichloropyrrole containing MS/MS fragment (*m/z* 161.9527 calcd for C_5_H_2_Cl_2_NO^+^, 161.9513, - 8.65 ppm, Figures S4-5). These data indicated that compound **2** is a dichlorinated analogue of **1**. To assign the positions of the two chlorines on the pyrrole, we recorded ^1^H, HSQC, and HMBC NMR spectra using a 600 MHz NMR instrument equipped with a 5 mm cryoprobe (Table S2). As expected, the ^1^H spectrum was almost identical with that of **1**, except for an additional sharp singlet at 7.04 ppm, the shielded of H-6 and H-14 resonances (from 3.65 to 3.50 and 3.88 to 3.49 ppm, respectively), and the slightly shielded H_3_-20 resonance from 0.99 to 0.92 ppm. In the HMBC spectrum of **2**, we observed a long-range correlation from the proton resonance at 7.04 ppm to the C-15 carbonyl, which established the structure of indanopyrrole B (**2**) as a 17- deschloro derivative of **1**.

### Pattern-based identification of the candidate indanopyrrole (*idp)* BGC

Based on biosynthetic precedent for the pyrrole containing natural products indanomycin,^27^ chlorizidine,^19^ armeniaspirol,^20^ and marinopyrrole,^21^ we anticipated that indanopyrrole biosynthesis is initiated using a proline derived di- or trichloropyrrole starter unit. As such, a DIAMOND BLAST^38^ database containing 42 MAR4 genomes was created using cblaster^39^ and queried for the proline adenyltransferase (*idmJ*) and prolyl-carrier protein dehydrogenase (*idmI*) gene sequences from the indanomycin (*idm*) BGC (MIBiG #: BGC0000079).^40^ This analysis returned 33 hits in 25 MAR4 genomes all within the bounds of BGCs called by AntiSMASH 5.0.^41^

Surprisingly, three distinct BGCs containing the pyrrole biosynthetic hooks were identified in the indanopyrrole producing strain CNX-425. To narrow down the candidates, we queried extracts of the same 42 strains for the indanopyrrole A molecular ion (*m/z* of 458.1060 ± 0.01) and detected it in seven. These seven strains shared one BGC with the pyrrole biosynthetic hooks, and it had the highest antiSMASH similarity score (39%) with *idm*. Using this pattern-based genome mining approach,^17^ we identified this conserved BGC as the top candidate for indanopyrrole biosynthesis and named it *idp* (Figure 1A, S6). In total, 15 of the 42 MAR4 genomes contained full-length *idp* BGCs, while five additional strains contained partial BGCs located on contig edges (Figure S6). Among the seven strains found to produce indanopyrroles, two contained partial BGCs suggesting that the truncations are artifacts of the sequencing and assembly process. While some strain-level variation among *idp* BGCs was observed (Figure S6), there is little evidence to suggest that some may be nonfunctional.

**Figure 1.**
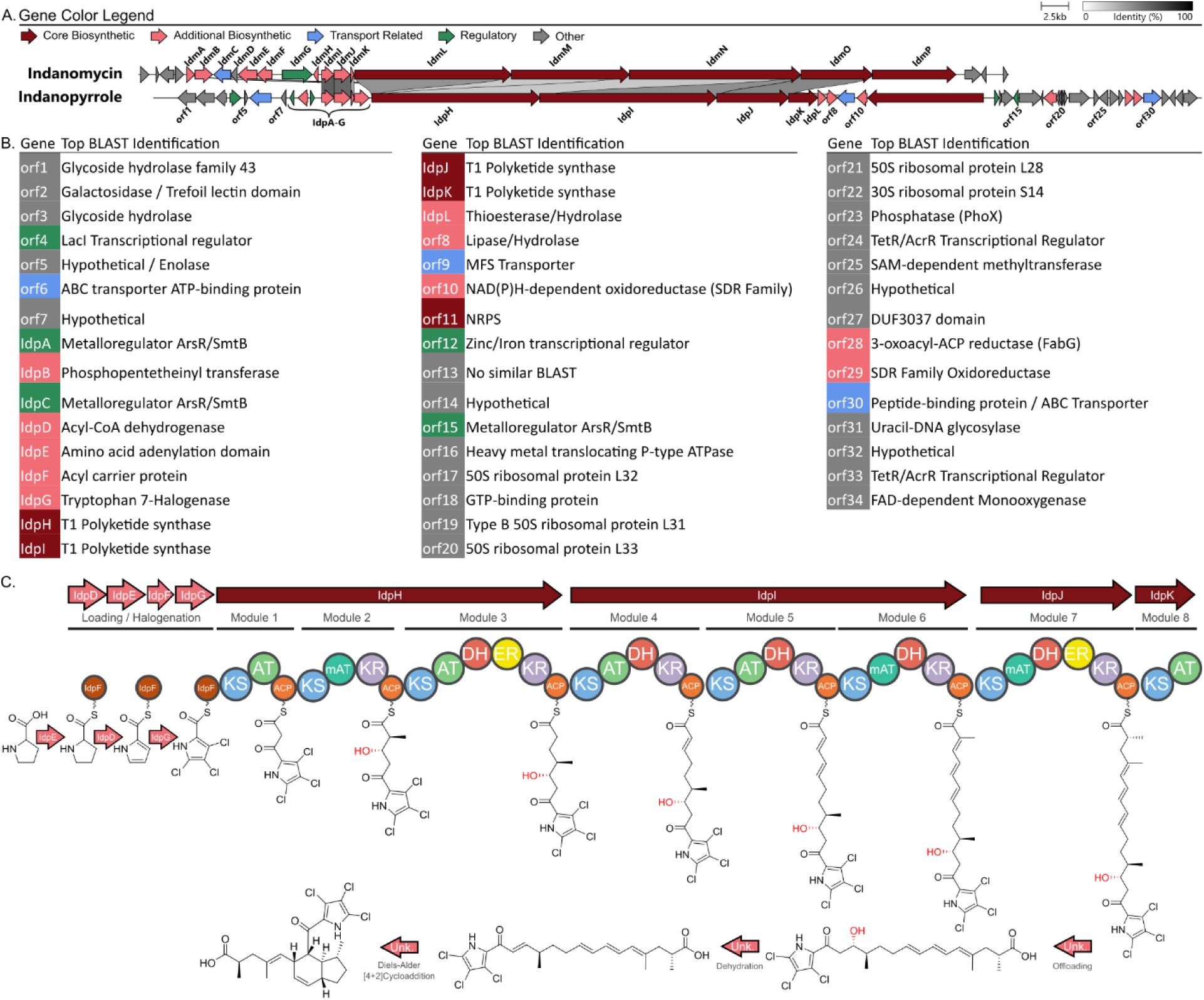
Candidate indanopyrrole BGC and proposed biosynthesis. A. Comparison of the candidate indanopyrrole BGC (*idp*) from *Streptomyces* sp. CNX-425 with the indanomycin BGC (*idm*, MIBiG #: BGC0000079). B. *Idp* gene annotations based on top BLAST matches to the NCBI NR/NT database. C. Proposed indanopyrrole biosynthesis in concordance with the *idp* BGC. Cryptic dehydration of the hydroxy group (in red) is predicted to afford an alkene dienophile intermediate facilitating an intramolecular Diels-Alder ([4+2] cycloaddition) reaction to form the indane.

Noting low production in CNX-425, we scaled-up the cultivation of additional *idp* containing MAR4 strains and found the highest production (5 mg indanopyrrole A and 0.5 mg indanopyrrole B from a 3 L culture) in strain CNY-716. Interestingly, indanopyrrole was not originally observed in this strain (Figure S6), but was instead observed in a scale up- fermentation following the addition of XAD-7 resin (Figure S7, see Methods). The additional compound obtained from this strain facilitated the structure elucidation and was used for expanded antibiotic screening.

### Biosynthesis of the indanopyrroles

A comparison of the *idp* and *idm* BGCs provides insight into the structural differences between their small molecule products (Figure 1A-B). As expected, both BGCs contain homologous genes encoding proline adenyltransferases (*idpE/idmJ*), proline carrier proteins (*idpF/idmK*), and dehydrogenases (*idpD/idmI*) to account for the generation of the pyrrole moieties from proline (Figure 1C). *Idp* then diverges by the presence of the halogenase gene *idpG*, which was annotated as a tryptophan halogenase. Unlike other BGCs linked to halogenated MAR4 natural products, *idp* does not contain a vanadium haloperoxidase encoding gene. Instead, *IdpG* shares homology with halogenase genes in the BGCs of other chlorinated pyrrole containing natural products^19,20,42^ and likely accounts for the tri- and dichlorinated pyrrole moieties observed in **1** and **2**, respectively. The chlorinated pyrrole generated from *idpD-G* then serves as the starter unit for seven polyketide extensions encoded by the three T1PKS genes (*idpH-J*). Notably, a NaPDoS2^43,44^ analysis of the module 1 (loading module) ketosynthase (KS) domain within *IdpH* places it in a clade with other pyrrole accepting KSs, further supporting the functional prediction for the starter unit (Figure S8).

Following starter unit selection, the acyltransferase (AT) domains associated with modules two, six, and seven are predicted to select for methylmalonyl-CoA. Based on conserved tryptophan and histidine residues, the ketoreductase (KR) domain within module 2 is assigned to the A2- type, which produce 2*S-*methyl-3*S-*hydroxy intermediates (Figure S9).^45^ Module 3 (*idpH*) contains the full suite of domains (KR, DH, and ER) to generate the alkane while modules 4-6 (*idpI*) contain KR and dehydratase (DH) domains that would afford a conjugated triene. Module 7 contains the full suite of domains to generate a branched alkane that, based on the lack of a conserved tyrosine residue in the active site of the *idpJ* enoylreductase (ER) domain, is predicted to install *R* configuration (Figure S10).^45^

The terminal T1PKS module (module 8, *idpK*) is comprised of a KS and AT domain and was observed in all *idp* BGCs except for one that appears to be truncated before this gene. This unusual domain organization resembles the terminal PKS module reported for *idm,*^27^ which differs only by the presence of a terminal cyclase domain. In both *idp* and *idm*, the AT domain within the terminal module lacks the active site residues required for selection and loading of the extender unit and is therefore predicted to be nonfunctional (Figure S11). However, *idp* further differs from *idm* in that the KS domain of the terminal module lacks the active site residues required for decarboxylative condensation and is therefore also predicted to be inactive (Figure S12). The inability of this module to support chain extension is supported by the structures of **1**-**2**. A stand-alone thioesterase (TE) domain (*idpL*) located immediately after *idpK* is homologous to *idmA*, which was presumed to be associated with chain release from the megasynthase during indanomycin biosynthesis.^46^ However, an analysis using the THYME thioesterase database of the *idpL* and *idmA* genes places them in the TE18 family of “editing” type II TEs, which remove prematurely decarboxylated extender units, stalled intermediates, or improperly edited CoA-bound starter units.^47,48^ Thus, it does not appear that *idpL* is involved in chain release. In the case of indanomycin, offloading has been proposed to involve the terminal cyclase domain within *idmP*,^27,49^ however the lack of this domain in *idpK* suggests either a different mechanism for **1**-**2** or that the cyclase is not involved (Figure S13). Finally, while antiSMASH calls a larger BGC, manual analyses have led us to propose that the *idp* BGC is best represented by *idpA-idpL*. As in indanomycin biosynthesis, questions remain about how the linear precursor predicted from the BGC is offloaded and cyclized to yield the final pyrroloketoindane products.

### Indane formation

Further comparison of the *idm* and *idp* BGCs and their products provides insight into the formation of the indane. In the case of indanomycin, it was suggested that *idmH* is an indane cyclase catalyzing a Diels-Alder [4+2] cycloaddition reaction. However, genetic knockout experiments lowered but did not abolish compound production^50^ and molecular modelling of the crystal structure did not reveal the potential for enzymatic activity.^51^ Notably, only one gene with similarity to *idmH* was detected among the indanopyrrole producing MAR4 strains (CNY-716, 36% amino acid similarity) and it was not located within any of the BGCs identified in that genome. Thus, the *idmH* mediated mechanism of indane ring formation proposed for indanomycin does not appear to apply to indanopyrrole. While several unannotated *idp* open reading frames could account for indane formation, it is intriguing to consider that the conserved KS domain associated with the non-elongating, terminal PKS module observed in *idm* and all *idp* gene clusters may be involved. KS functional diversification is well documented and includes the non-elongating, terminal KS domain in salinosporamide A biosynthesis, which was recently shown to catalyze the formation of an alkene which then facilitates intramolecular carbon-carbon bond formation and the cyclized end product.^52^ Notably, both *idm* and *idp* lack a dehydratase domain in the second PKS module, which would provide the alkene dienophile required for indane ring formation via a Diels-Alder [4+2] cycloaddition reaction.^27,53–56^ It is intriguing to speculate that the terminal, non-elongating, KS domains observed in both *idm* and *idp* catalyze this cryptic dehydration step.

To address the potential neofunctionalization of the *idpK* KS domain, we modelled the 3D protein structure using the open-source implementation of AlphaFold 2^57^ in ColabFold.^58^ The model showed strong structural homology with the crystal structure of the KS domain from 6- deoxyerythronolide B synthase (DEBS), with the active site residues collocated within the substrate binding pocket (Figure S14A). Molecular docking studies using the AMDock suite^59^ showed that indanopyrrole A (**1**) fits within the predicted substrate binding pocket of the *IdpK* KS domain. In the model, the carboxylic acid moiety of **1** is adjacent to the active site cysteine (Cys-167). This position is homologous to DEBS Cys-211, which forms a thioester bond with the polyketide intermediate generated by the preceding PKS module (Figure S14A inset). This spatial orientation aligns active site histidine-302 above C-14 in the indanopyrrole structure. Positioning this residue above the carbon alpha to the carbonyl group could allow the active site histidine to abstract a hydrogen and initiate a cascade reaction that ends with the elimination of water from the C-13 hydroxy group thus accounting for the cryptic dehydration step and the formation of the alkene dienophile (Figure S14B). Finally, the shape of the substrate binding pocket conforms to the shape of indanopyrrole A, suggesting it could promote a conformational change in the linear precursor that facilitates a spontaneous Diels-Alder reaction following dehydration (Figure S14A, inset). However, we were unable to model the proposed linear precursor within the active site to address this hypothesis. It should be noted that the absolute configuration of the indane ring systems in **1** and **2** are opposite from the indanomycins, which could account for the low KS domain sequence similarity and the differences in active site residues. While speculative, assigning dehydratase activity to the terminal KS domains in *idm* and *idp* would indicate a new mechanism for indane formation in the indanopyrroles and other pyrroloketoindanes.

### Gene cluster and metabolite distribution

To more broadly explore *idp* and *idm* distributions, we used cblaster^60^ to remotely query the NCBI nr database using the respective adenyltransferase (*idpE/idmJ)* and proline dehydrogenase (*idpD/idmJ)* genes, which returned 3,283 (*idp*) and 3,198 (*idm*) hits. After filtering to include only those that contained at least one gene with >70% homology to a PKS gene within *idp* or *idm* and duplicate removal, 40 BGCs were identified and manually assigned to seven BGC groups and nine singletons based on gene synteny (Figure S15). AntiSMASH analyses revealed top matches to the indanomycin, nargenicin, and calcimycin BGCs along with three with no known product (Figure S15). Three complete *idm* BGCs were identified in *Streptomyces albireticuli* strains, NRRL B1670 (JAJQQQ010000001.1), NRRL B1670 Type B (JAJQQR010000001.1), and NRRL B1670 Type C (JAJQQS010000003.1) (Figure 2), all of which lacked the terminal cyclase domain observed in *idmP* (Figure S13). Additionally, two fragmented *idm* BGCs were detected on contig edges in *S. ureilyticus* (NZ_JAAKZX010000175.1) and *S. coffeae* (NZ_JAERRF010000046.1). One putative full-length *idp* BGC was observed in *Micromonospora* sp. WMMC 250 (NZ_JAPZBJ010000002.1) (Figure 2) and one fragmented BGC was identified on a contig edge in the genome of *Streptomyces sedi* JCM 16909 (VDGT01000017.1). These results indicate that, despite being detected in genetically distant actinobacterial lineages (*Streptomyces* and *Micromonospora*), both *idm* and *idp* are rare in publicly available sequence data.

**Figure 2.**
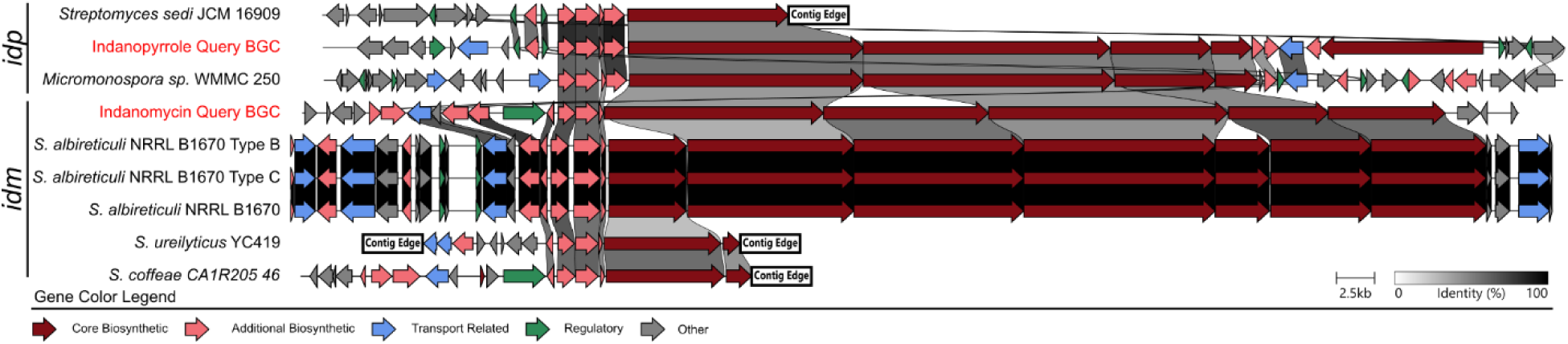
Distribution of *idp* and *idm*. The seven closest *idp* and *idm* homologs identified from GenBank searches highlight the rarity of these BGCs and presents opportunities for discovery. A full list of BGC hits is provided in Supplementary Figure S15.

We next used the Mass Spectrometry Search Tool^61^ (MASST) within the Global Natural Products Social Molecular Networking^62^ (GNPS) platform to search for the MS/MS spectrum of indanopyrrole A within public datasets. After detecting no hits using the default parameters, we lowered the minimum cosine score to 0.6 and the minimum matched fragments to 2 and detected 71 hits when allowing for analogue searching. Upon manual inspection, none of the *m/z* values or fragmentation spectra matched indanopyrrole A. The lack of any significant matches among 2,709 public datasets highlights the rarity of these natural products.

### Bioactivity

Indanopyrrole A demonstrated significant antibacterial activity, with a minimum inhibitory concentration (MIC) of 4 µg/mL against membrane-deficient *Escherichia coli* lptD4213^63^ (Table 1). Broader testing revealed potent activity against several clinically relevant Gram-positive pathogens, including methicillin-resistant *Staphylococcus aureus* TCH1516 (MIC = 2 µg/mL), group A *Streptococcus* M1T1 (MIC = 4 µg/mL), vancomycin-resistant *Enterococcus faecium* DAPS (MIC = 2 µg/mL), and methicillin-resistant *Staphylococcus epidermis* (MIC = 4 µg/mL) (Table 1). To our best knowledge, **1** is the first pyrroloketoindane natural product to show activity against Gram-negative organisms, showing MIC values of 1-2 µg/mL against *Haemophilus influenzae* (Table 1). In contrast, indanopyrrole B (**2**) was inactive at all concentrations tested, likely due to differences in chlorination between the two compounds. The cytotoxicity of **1** was measured at 16 µg/mL after 24 hours of exposure in an A549 cell line viability assay (Figure S16), indicating an antibiotic therapeutic index of 4-8 (ratio of antibiotic to cytotoxic activity).

**Table 1.** Antibacterial activities of indanopyrroles A and B reported as minimum inhibitory activity. N.T. = Not Tested.

| Strains tested | Gram +/- | Indanopyrrole A | Indanopyrrole B |
| --- | --- | --- | --- |
| MRSA TCH1516 | + | 2 µg/mL | >16 µg/mL |
| Group A *Streptococcus* M1T1 | + | 4 µg/mL | >16 µg/mL |
| Vancomycin-resistant *Enterococcus faecium* DAPS | + | 2 µg/mL | >16 µg/mL |
| MRSE (*Staphylococcus epidermidis*) | + | 4 µg/mL | N.T. |
| *Haemophilus influenzae* 1 | - | 1 µg/mL | N.T. |
| *Haemophilus influenzae* 2 | - | 2 µg/mL | N.T. |
| *Escherichia coli* LptD4213 | - | 4 µg/mL | N.T. |
| *Escherichia coli* K1 RS218 | - | >32 µg/mL | >16 µg/mL |
| *Acinetobacter baumannii* 5075 | - | >32 µg/mL | N.T. |

Given the antibiotic activity of indanopyrrole A, it is unclear how producing strains maintain resistance. However, a drug resistance transporter related to the *Burkholderia thailandensis EmrB* multidrug efflux pump is conserved in all *idp* BGCs suggesting a possible resistance mechanism. Similar mechanisms of drug resistance in Gram-negative organisms such as *Acinetobacter baumanii* and *Escherichia coli* are mediated by upregulation or genomic expansion of efflux pumps and may explain the lack of activity against these strains.^64,65^

### Conclusions

The MAR4 group of marine actinomycetes has been a prolific source of halogenated natural products.^6^ To further explore their biosynthetic potential, we applied pattern-based genome mining to 42 strains, focusing on identifying new halogenated metabolites and their candidate BGCs. This effort led to the discovery of the pyrroloketoindane natural products indanopyrroles A and B (**1**-**2**), which are the seventh and eighth pyrroloketoindane natural products described and the only halogenated members of this rare compound class. Comprehensive 1D and 2D NMR spectroscopy facilitated the structure elucidation of **1**-**2** including their relative configuration, while their absolute configuration was proposed based on bioinformatic prediction of the stereoselective KR and ER domains in the candidate BGC.

Indanopyrrole production was linked to a BGC (*idp*) containing genes associated with chloropyrrole biosynthesis. As in the biosynthesis of the related compound indanomycin, a cryptic dehydration step is predicted to facilitate an intramolecular Diels-Alder cycloaddition that yields the final cyclized product. Comparisons between the *idp* and *idm* BGCs suggest that the terminal, non-elongating KS domains observed in these BGCs could account for this cryptic dehydration. While *in-silico* docking studies support this hypothesis, experimental verification is still needed. A search of the NCBI nr database revealed seven BGCs with similarity to *idp* or *idm*, supporting the rarity of pyrroloketoindane natural products. All seven of these BGCs, present in diverse actinomycetes, contain terminal KS domains homologous to *idpK* and *idmP*, supporting a functional role for this domain in the pathway.

Indanopyrrole A (**1**) exhibits potent, broad-spectrum antibiotic activity and, to the best of our knowledge, is the first compound in this class with reported activity against Gram-negative bacteria. The inactivity of indanopyrrole B (**2**) emphasizes the importance of the tri-chlorinated pyrrole moiety for the activity of **1**. Unlike the pyrroloketoindane antibiotic indanomycin, which acts as an ionophore for divalent cations, **1** lacks the tetrahydrofuran moiety known to coordinate metals^27^ indicating it may function via a different mechanism. The promising therapeutic index of indanopyrrole A (**1**) supports further structure-activity studies and antibiotic lead development.

This work showcases the value of pattern-based genome mining for discovering new antibiotics and their BGCs within the biosynthetically gifted MAR4 group of marine bacteria. It also expands the breadth of halogenated natural products reported from the MAR4 actinomyces and provides new insights into potential KS functional diversification and indane ring formation in the rare pyrroloketoindane compound class.

## EXPERIMENTAL SECTION

### General experimental procedures

Optical rotations were recorded on a Jasco P-2000 polarimeter. UV spectra were measured on a Jasco V-630 spectrophotometer. IR spectra were acquired on a JASCO FTIR-4100 spectrometer (Jasco Corp., Tokyo, Japan). NMR spectra (1D and 2D) were measured at 23°C on a JEOL ECZ spectrometer (500 MHz) equipped with a 3 mm ^1^H{^13^C} room temperature probe (JEOL, Akishima, Tokyo, Japan) or on a Bruker Avance III (600 MHz) NMR spectrometer with a 5 mm ^1^H{^13^C/^15^N} room temperature or cryo probe (Billerica, MA). ^13^C NMR spectrum was recorded on a Varian 500 MHz spectrometer equipped with a 5 mm ^1^H{^13^C} XSens cold probe (Varian Inc., Palo Alto, CA, USA, now Agilent Technologies). NMR spectra were referenced to the solvent signals (CHD_2_OD, δ_H_ 3.31 and CD_3_OD, δ_C_ 49.00 ppm). LC- HR-ESIMS was performed on an Agilent 1260 Infinity HPLC system equipped with a degasser, binary pump, autosampler, DAD detector, coupled to an Agilent 6530 Accurate-Mass QToF with ESI-source coupled with an Agilent 1260 Infinity HPLC and calibrated using the Agilent Reference Calibration Mix. Compounds were isolated on an Agilent HPLC system with 1100 G1312A binary pump, 1100 G1315A DAD UV/Vis detector, 1100 G1313A autosampler, and 1100 G1322A degasser (Agilent Technologies, Santa Clara, CA).

### Small-scale strain cultivation and metabolite extraction

All cultures were grown at 28°C and shaken at 230 rpm with metal springs in A1 media containing potassium bromide (10 g/L starch, 4 g/L yeast extract, 2 g/L peptone, 22 g/L instant ocean, 0.1 g/L KBr). Cryogenic stocks of 40 MAR4 strains as well as lyophilized material for the two MAR4 type strains purchased from the DSMZ (DSM 41644, DSM 41902) were inoculated as a preculture into 50 mL of media. After 7 days, 0.5 mL was transferred into 50 mL of the same medium for a second seed culture. After 5 days, 5 mL of the second seed culture was frozen for DNA analysis and 0.5 mL was used to inoculate 50 mL of fresh media. After four days, HP-20 resin (1 g, wet weight) was added to each flask. After three days of incubation with resin, all cultures were extracted with 50 mL EtOAc. Organic extracts were separated, dried with anhydrous Na_2_SO_4_, filtered, concentrated by rotary evaporation, and stored at -20°C until further analysis.

### Large-scale cultivation and extraction of strain CNY-716

Aliquots (10 mL) of a 50 mL preculture of *Streptomyces* sp. CNY-716 were inoculated into three 2.8 L Fernbach flasks containing 1 L of A1 medium and incubated at 28°C and 120 rpm. On day 10, activated sterile XAD-7 resin (20 g, ThermoFisher Scientific) was added to each flask and the cultures incubated for an additional 4 days. After 14 days, the resin and the cellular material were filtered through cheesecloth, washed with deionized H_2_O, and extracted with MeOH (4 x 200 mL). The solvent was removed from the pooled extracts under reduced pressure to yield a black oily material (2.0 g).

### Fractionation and Isolation

The CNY-716 crude extract was fractionated using vacuum liquid chromatography (15 g C_18_ silica gel) and a step gradient of MeOH:H_2_O (50 mL each; 25:75, 50:50, 60:40, 70:30, 80:20, 90:10, and 2x 100:0) into eight fractions (LC1-8). Fractions containing indanopyrroles (LC6-7) were further purified using HPLC (Phenomenex Kinetex C_18,_ 5 µm, 150x4.6 mm column, isocratic ACN:H_2_O 68:32 with 0.05% formic acid mobile phase, 1.3 mL/min flow rate) to yield 5 mg of indanopyrrole A (**1**, retention time t_R_ = 12 min) and 0.5 mg indanopyrrole B (**2**, t_R_ = 6 min).

### Compound characterization

*Indanopyrrole A (**1**)*: amorphous white solid; [α]^22^ –214 (*c* 0.26, MeOH); UV (MeOH) λ_max_ (log ε) 200 (3.59), 261 (sh) (2.97) 295 ((3.34); IR (neat) ν_max_ 3186 cm^-1^, 2938 cm^-1^,2866 cm^-1^, 1707 cm^-1^, 1639 cm^-1^, 1445 cm^-1^, 1396 cm^-1^, 1372 cm^-1^, 1226 cm^-1^, 1018 cm^-1^; ^1^H, ^13^C and 2D NMR, Table S1; HR-ESI-TOF-MS *m/z* 458.1060 [M+H]^+^ (calcd for C_22_H_27_Cl_3_NO_3_^+^, 458.1052, 1.75 ppm).

*Indanopyrrole B (**2**)*: amorphous white solid; [α]^22^ -111 (*c* 0.26, MeOH); UV (MeOH) λ_max_ (log ε) 200 (3.51), 245 (3.01) 296 (3.51); IR (neat) ν_max_ 3326 cm^-1^, 2952 cm^-1^,2866 cm^-1^, 1644 cm^-1^, 1410 cm^-1^, 1027 cm^-1^; ^1^H, ^13^C and 2D NMR, Table S2; HR-ESI-TOF-MS *m/z* [M+H]^+^ 424.1451 (calcd for C_22_H_28_Cl_2_NO_3_^+^, 424.1441, 2.36 ppm).

### Genomic DNA Extraction

DNA was extracted from frozen aliquots (5 mL) of the 42 MAR4 strains using the Promega Wizard Genomic DNA Purification Kit with suggested modifications for Gram-positive bacteria. DNA purity, concentrations, and size were assessed using NanoDrop, Qubit, and gel electrophoresis. Short-read, paired-end Illumina sequencing (PE150) was performed at SeqCenter (Pittsburgh, PA). Initial genome assembly was performed by quality filtering raw reads with the BBMap Toolkit^66^ followed by a preliminary assembly using SPAdes.^67^ All assemblies were compared for whole-genome average nucleotide identity (ANI) using fastANI^68^ to identify strains sharing 95% ANI. Ten representative strains from each 95% ANI clade were selected for long-read Nanopore sequencing (Oxford) and compiled with public SRA data from previously sequenced strains (N=12 Illumina, N=3 PACBIO). Data were combined for each strain to perform a hybrid assembly with unicycler^69^ using a kmer count 31,41,51,61,71,81,91,95,101,105,111 and “mode” determined by ANI similarity to a reference strain (i.e., “bold” with ANI=100%, “normal” with ANI>99%, “conservative” with ANI >97%). All assemblies were checked for quality, completeness, and contamination with checkM.^70^ Biosynthetic gene clusters (BGCs) were predicted using antiSMASH v5^41^ and clustered into gene cluster families using BiG-SCAPE^71^ with GCFs defined at 0.4 dissimilarity based on a combined metric of gene synteny, protein domain structure, and homology. MAR4 BGCs were annotated using the MIBiG 2.7 database.^72^ Network files from BIG-SCAPE were visualized in Cytoscape 3.10.^73^ Synteny plots were generated from GBK files created by AntiSMASH using the clinker package with default parameters.

BGC amino acid sequences were sourced from NCBI GenBank and the AntiSMASH output. For each gene alignment Muscle5^74^ was used to create a stratified ensemble of 16 alignments using the default parameters. The alignment with the highest column confidence value was extracted from the ensemble using the maxcc option and used for comparison of active site residues.

### Metabolomics and Mass Spectrometry

Dried crude extracts were resuspended in MeOH (1 mg/mL) and centrifuge filtered using 0.2 µm filters (American Chromatography Supply, Vineland NJ). Samples (5 µL) were injected into an Agilent 1290 HPLC coupled to an Agilent 6530 quadrupole time-of-fight (QToF) spectrometer. Parameters were set to a flow rate of 0.75 mL/min through a Kinetex C_18_ reversed-phase column (5 μm, 150x4.6 mm) under the following conditions: 0-4 min 5% acetonitrile (0.1% TFA) in water (0.1% TFA) with this first 4 minutes diverted to waste, 4-34 min: 10-100% acetonitrile (0.1% TFA) in water (0.1% TFA), 34-36 min 100% acetonitrile, 36-36.5 min 100-5% acetonitrile, 36.5-40 min 5% acetonitrile. MS1 data was collected in positive and negative over two runs with a mass range of 80-1700 *m/z* acquiring three spectra per second, MS2 fragmentation data were collected using two scans per second with a collision energy of 30eV. The source gas temperature was 300 °C at a flow rate of 11 L per minute at 35 psig.

### Molecular modelling and docking studies

A 3D model of indanopyrrole A (**1**) was built in Spartan’24 V1.1.0 and minimal energy conformers searched using molecular mechanics (CorrMMFF, ΔE <25 kJ/mol) and further optimized using DFT (Est. Density Functional ωB97X- D/6-31G*). The online version of ColabFold (https://colab.research.google.com/github/sokrypton/ColabFold/blob/main/AlphaFold2.ipynb) was used to generate a 3D structure from the amino acid sequence of the *IdpK* KS domain. Default parameters were used except the “template_mode” was changed to include the pdb70 reference library. Both 3D models were used as input for docking analysis using AMDock.^75^ The default search parameters were used in an Autodock Vina model.^76,77^

### Antibacterial testing

**1** was ten-fold serially diluted (from 128 µg/mL to 0.25 µg/mL) in a 96- well plate using sterile medium (Muller-Hinton broth; 50 µL of each solution per well). An *E. coli* LptD 4213 inoculum (50 µL) was added to reach a final concentration of 2e5 CFU/mL (determined via OD_600_). The final test concentrations ranged from 64 µg/mL to 0.125 µg/mL. After incubation for 18 hours at 37 °C, the well with the lowest concentration of compound that did not exhibit microbial growth was determined to be the MIC.

Additional MIC values were determined using broth microdilution in accordance with the Clinical Laboratory Standards Institute (CLSI) guidelines using cation-adjusted Mueller Hinton Broth (MHB) with minor modifications. Briefly, bacteria were grown to mid-log phase (OD_600nm_ = 0.4) at 37 °C while shaking except for GAS which was grown under static condition. Bacterial cells were then centrifuged, washed, and diluted in PBS to obtain 2×10^6^ cfu/mL with 10 μL added to individual wells of a 96-well plate containing 170 μL MHB. Serial dilutions of indanopyrroles A and B starting at 32 μg/mL or 16 μg/mL, respectively, were made in a separate plate, 20 μL of the compound was then added to the test plate. The plates were sealed with parafilm and incubated at 37 °C for 24 h. Turbidity was measured at OD_600nm_ using an EnSpire Alpha plate reader. MIC was defined as the lowest concentration of the test compounds that inhibited bacterial growth.

### Cell Viability Assay

A549 cells were seeded in 24 well plates (Corning, United States) at 2x10^5^ cells/well. Cells were left untreated or treated with 16, 32, and 128 μg/mL of indanopyrrole A, or with corresponding amounts of the H2O:DMSO (1:2) solvent vehicle as a negative control. As a positive control, A549 cells were lysed with Triton X-100. Cell culture supernatants were collected at two time points: 2 and 24 hrs. Cellular cytotoxicity was assessed by measuring the levels of lactate dehydrogenase (LDH; Promega, United States) released by the host cell into supernatant. The percentage of cell death was calculated after subtracting the levels found in untreated control cells and dividing by the levels in a positive control of cells treated with a lysis solution (Triton X-100).

## ASSOCIATED CONTENT

### Supporting Information

Supplemental figures including MS and NMR spectra for all compounds, Genetic analyses, and 3D modeling (PDF)

## ACKNOWLEDGEMENT

The authors acknowledge assistance from Tadeusz Molinski (UCSD Chemistry and Biochemistry Department, Skaggs School of Pharmacy) for providing access to optical rotation, UV and IR instruments and expert guidance, as well as Brendan Duggan (Skaggs School of Pharmacy) and Anthony Mrse (Department of Chemistry, UCSD) for expert guidance and help in the NMR recording. This work was supported by the National Institutes of Health grant R01GM085770 to PRJ.

## Notes

### Competing Interest Statement

The authors have declared no competing interest.

