## Supplemental Material for "Pattern-based genome mining guides discovery of the antibiotic indanopyrrole A from a marine streptomycete"

*These authors contributed equally

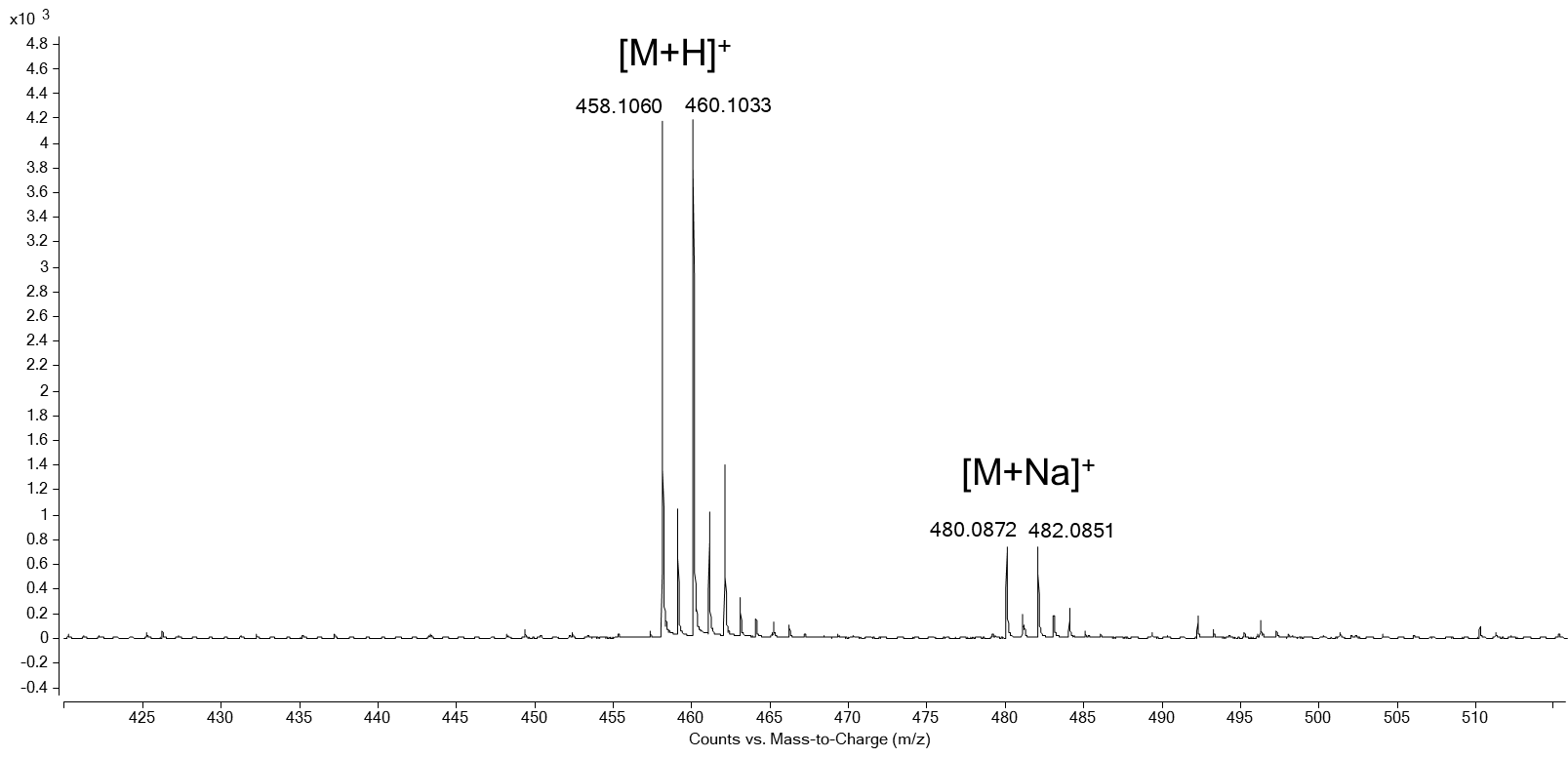

**Figure S1.** MS spectrum of indanopyrrole A (**1**), MF C_22_H_26_Cl_3_NO_3_.

**Table S1.** NMR data of indanopyrrole A (**1**).

| 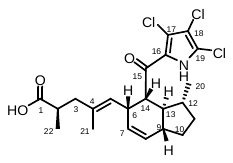 | | | | | |
| --- | --- | --- | --- | --- | --- |
| *^a^* | δ_C_ *^b^* | δ_H_ (*J* in Hz) | COSY | HMBC | NOESY |
| 1 | 181.5, C |  |  |  |  |
| 2 | 39.6, CH | 2.44, ddd (9.0, 6.8, 5.7) | H_2_-3, H_3_-22 | C-1, C-3, C-4, C-22 |  |
| 3 | 44.7, CH_2_ | a 1.89, dd (13.7, 9.0)  b 2.28, dd (13.7, 5.7) | H-2, H-5 (w)  H-2, H-5 (w) | C-1, C-2, C-4, C-5, C-22  C-1, C-2, C-4, C-5, C-22 |  |
| 4 | 133.7, C |  |  |  |  |
| 5 | 128.3, CH | 5.06, d (10.3) | H-3b, H-6, H_3_-21 | C-3, C-6, C-21 | H-3a, H-6, H-13 |
| 6 | 38.7, CH | 3.65, m | H-5, H-7, H-14 | C-7 | H-5, H-14, H_3_-21 |
| 7 | 130.0, CH | 5.33, dt (9.7, 3.4) | H-6, H-8, H-9 (w) | C-6, C-9, C-14 |  |
| 8 | 130.4, CH | 5.84, dt (9.7, 1.8) | H-7 | C-6, C-7, C-9, C-10, |  |
| 9 | 46.7, CH | 2.10, m | H-7 (w), H_2_-10 |  | H-10b, H-12,  H-14, |
| 10 | 28.6, CH_2_ | a 1.29 m  b 1.81, ddd (16.3, 9.2, 2.1) | H-9, H_2_-11  H-9, H_2_-11 | C-9, C-11,  C-12, C-13 |  |
| 11 | 34.4, CH_2_ | a 1.34, m  b 2.01, ddd (16.9, 12.8, 8.8) | H_2_-10, H-12  H_2_-10, H-12 | C-20  C-13 |  |
| 12 | 39.3, CH | 1.61, m | H_2_-11, H-13 H_3_-20 | C-11, C-13 |  |
| 13 | 47.8, CH | 1.58, m | H-9, H-12, H-14 | C-9, C-12, C-14, C-20 | H-5, H_3_-20 |
| 14 | 53.6, CH | 3.88, dd (10.5, 6.7) | H-6, H-13 | C-6, C-13, C-15, C-16 | H-6, H-9, H-12 |
| 15 | 189.8, C |  |  |  |  |
| 16 | 128.0, C |  |  |  |  |
| 17 | 116.1, C |  |  |  |  |
| 18 | 111.5, C |  |  |  |  |
| 19 | 120.6, C |  |  |  |  |
| 20 | 21.7, CH_3_ | 0.99, d (6.1) | H-12 | C-11, C-12, C-13 | H-13 |
| 21 | 15.8, CH_3_ | 1.31, br s | H-5 | C-3, C-4, C-5 | H-2, H-6, H-7 |
| 22 | 16.8, CH_3_ | 0.98, d (6.8) | H-2 | C-1, C-2, C-3 |  |

*^a^* All assignments are based on extensive 1D and 2D NMR measurements (COSY, HSQC, HMBC).

*^b^* Multiplicities determined by HSQC.

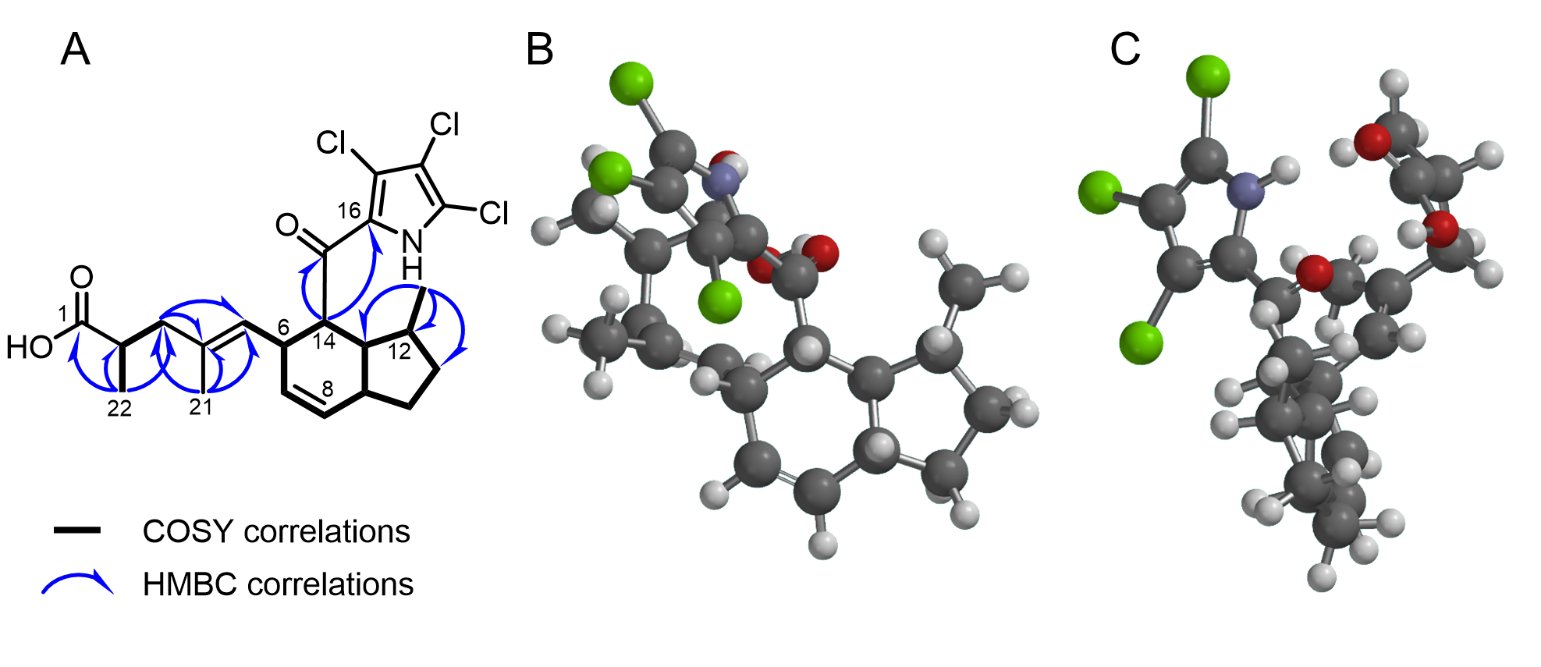

**Figure S2.** Structure elucidation of indanopyrrole A (**1**). A. COSY (bold bonds) and key HMBC (blue arrows) correlations. B-C. 3D model of minimal energy optimized conformer (Est. Density Functional ωB97X-D/6-31G*, 200.51 kJ/mol, 96.6% Boltzmann distribution).

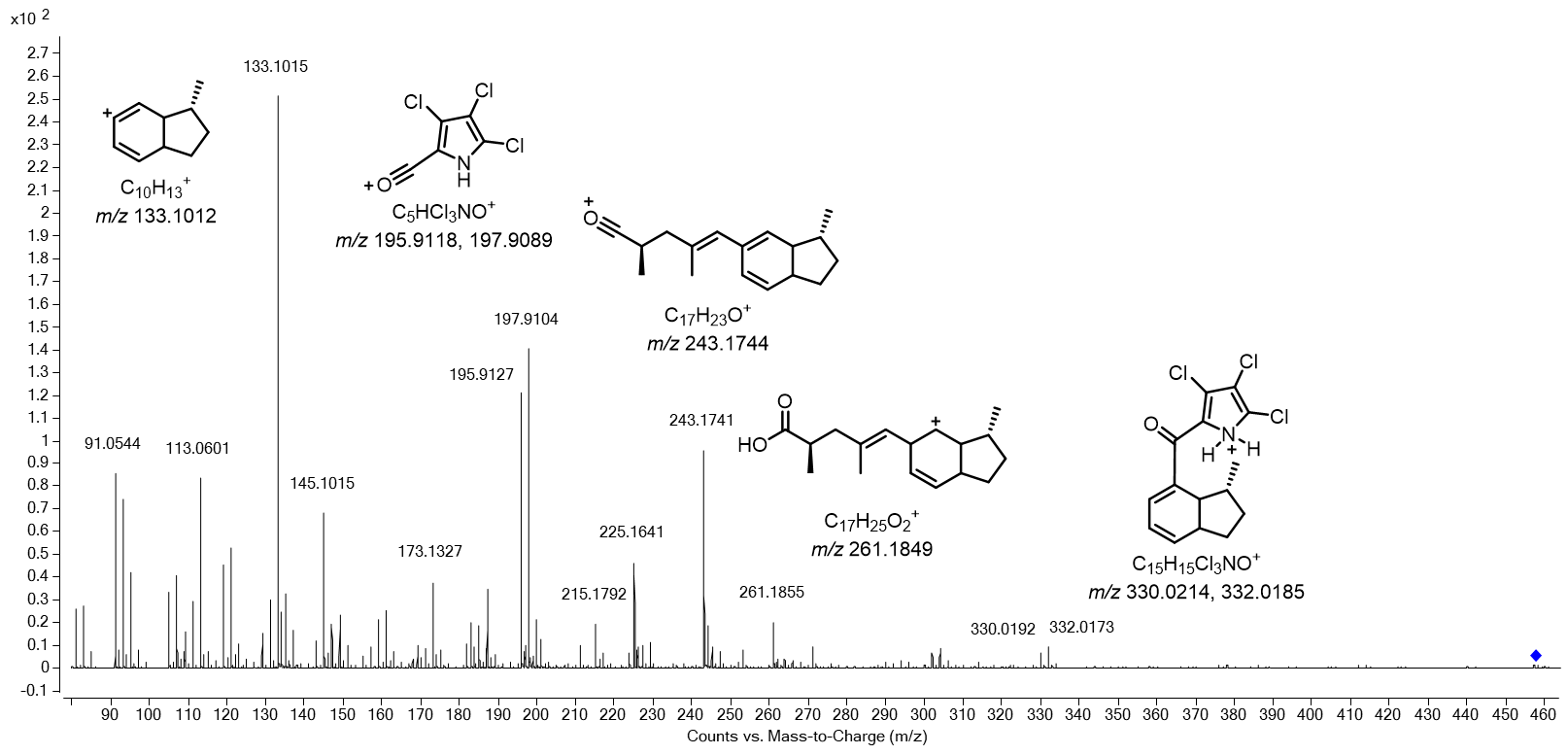

**Figure S3.** Annotated MS/MS spectrum of indanopyrrole A (**1**).

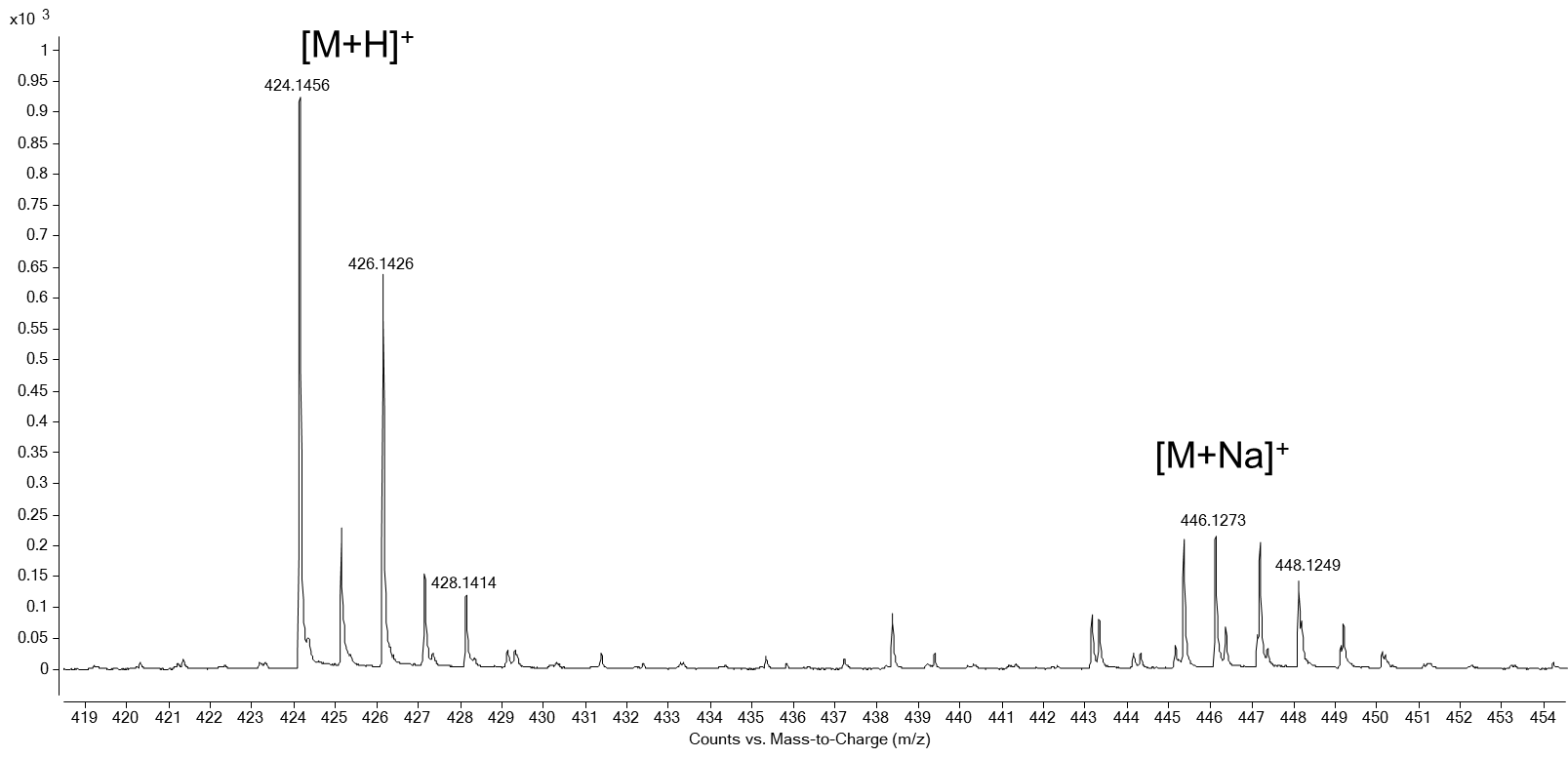

**Figure S4.** MS spectrum of indanopyrrole B (**2**) MF C_22_H_27_Cl_2_NO_3_.

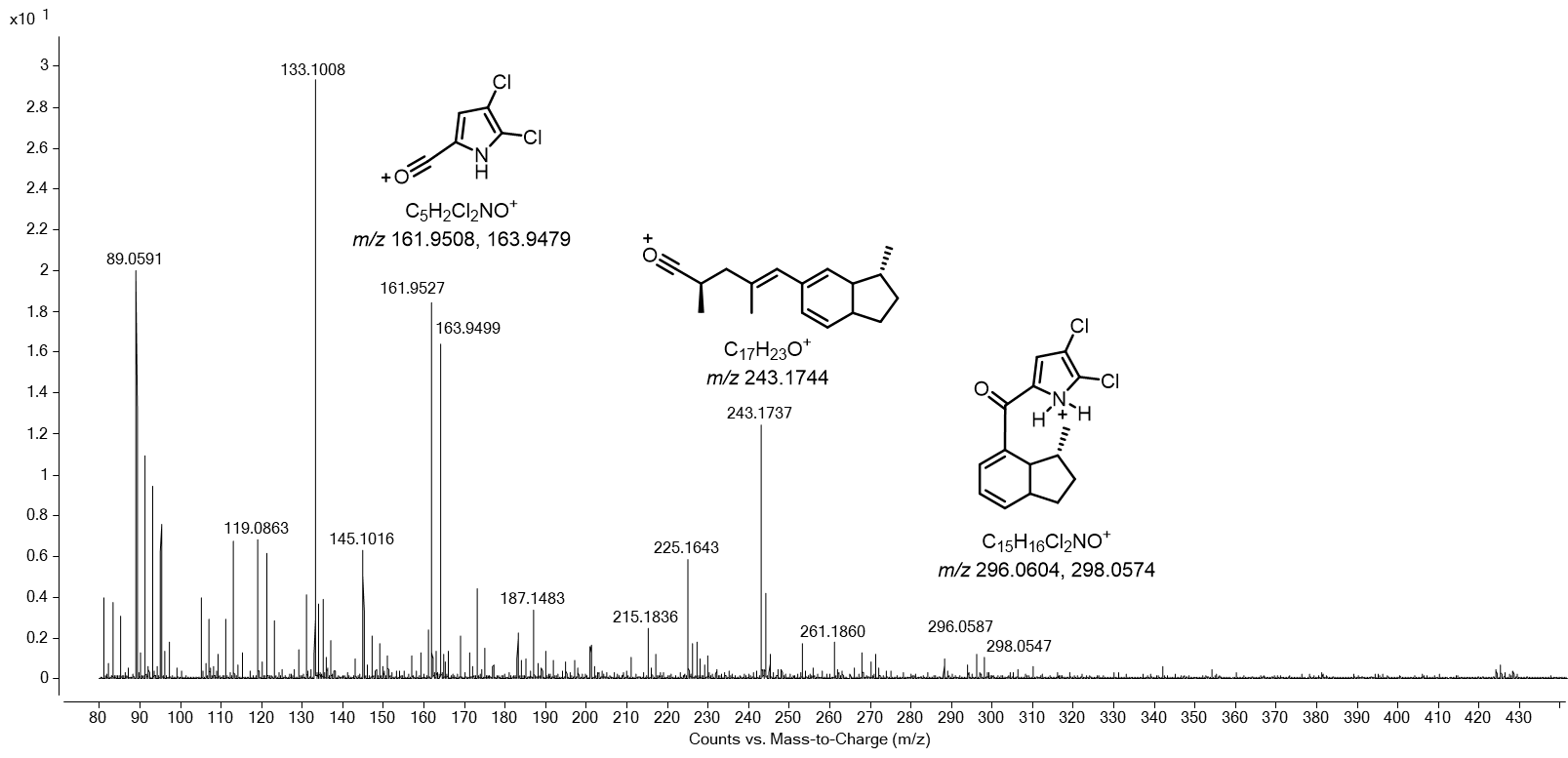

**Figure S5.** Annotated MS/MS spectrum of indanopyrrole B (**2**).

**Table S2.** NMR data of indanopyrrole B (**2**).

| 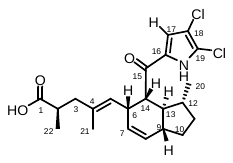 | | | | |
| --- | --- | --- | --- | --- |
| *^a^* | δ_C_ *^b^* | δ_H_ (*J* in Hz) | COSY | HMBC |
| 1 | 181.3, C |  |  |  |
| 2 | 39.2, CH | 2.45, ddd (5.4, 6.8, 9.1) | H_2_-3, H_3_-22 | C-1, C-3, C-4, C-22 |
| 3 | 44.7, CH_2_ | a 1.92, dd (9.1, 13.4)  b 2.29, dd (5.4, 13.4) | H-2, H-5 (w)  H-2, H-5 (w) | C-1, C-2, C-4, C-5, C-21, C-22  C-1, C-2, C-4, C-5, C-21, C-22 |
| 4 | 133.1, C |  |  |  |
| 5 | 128.2, CH | 5.10, br d (9.3) | H-3b, H-6, H_3_-21 | C-3, C-6 (w), C-21 |
| 6 | 40.2, CH | 3.50 m | H-5, H-7, H-14 | C-5 |
| 7 | 129.9, CH | 5.33, dt (3.0, 3.2, 9.8) | H-6, H-8, H-9 (w) | C-6 (w), C-9 |
| 8 | 130.0, CH | 5.83, br d (9.8) | H-7 | C-6, C-9, C-10, C-13 |
| 9 | 46.1, CH | 2.12, m | H-7 (w), H_2_-10 | C-8 (w) |
| 10 | 27.9, CH_2_ | a 1.27, m  b 1.81, m | H-9, H_2_-11  H-9, H_2_-11 | C-8, C-9, C-11  C-9, C-12, C-13 |
| 11 | 34.4, CH_2_ | a 1.32, m  b 2.01, m | H_2_-10, H-12  H_2_-10, H-12 | C-10, C-20  C-10, C-13 |
| 12 | 38.9, CH | 1.59, m | H_2_-11, H-13 H_3_-20 | C-11, C-13 |
| 13 | 47.7, CH | 1.51, dt (10.4, 10.6) | H-9, H-12, H-14 | C-6, C-9, C-12, C-14, C-20 |
| 14 | 53.2, CH | 3.49, m | H-6, H-13 | C-6, C-13, C-15, C-16 |
| 15 | 189.8, C |  |  |  |
| 16 | 131.5, C |  |  |  |
| 17 | 115.8, CH | 7.04, s |  | C-15, C-16, C-19 |
| 18 | n. o. |  |  |  |
| 19 | 121.2, C |  |  |  |
| 20 | 20.8, CH_3_ | 0.92, d (6.6) | H-12 | C-11, C-12, C-13 |
| 21 | 15.5, CH_3_ | 1.33, br s | H-5 | C-3, C-4, C-5 |
| 22 | 16.2, CH_3_ | 1.00, d (6.8) | H-2 | C-1, C-2, C-3 |

*^a^* All assignments are based on extensive 1D and 2D NMR measurements (COSY, HSQC, HMBC).

*^b^* Multiplicities determined by HSQC.

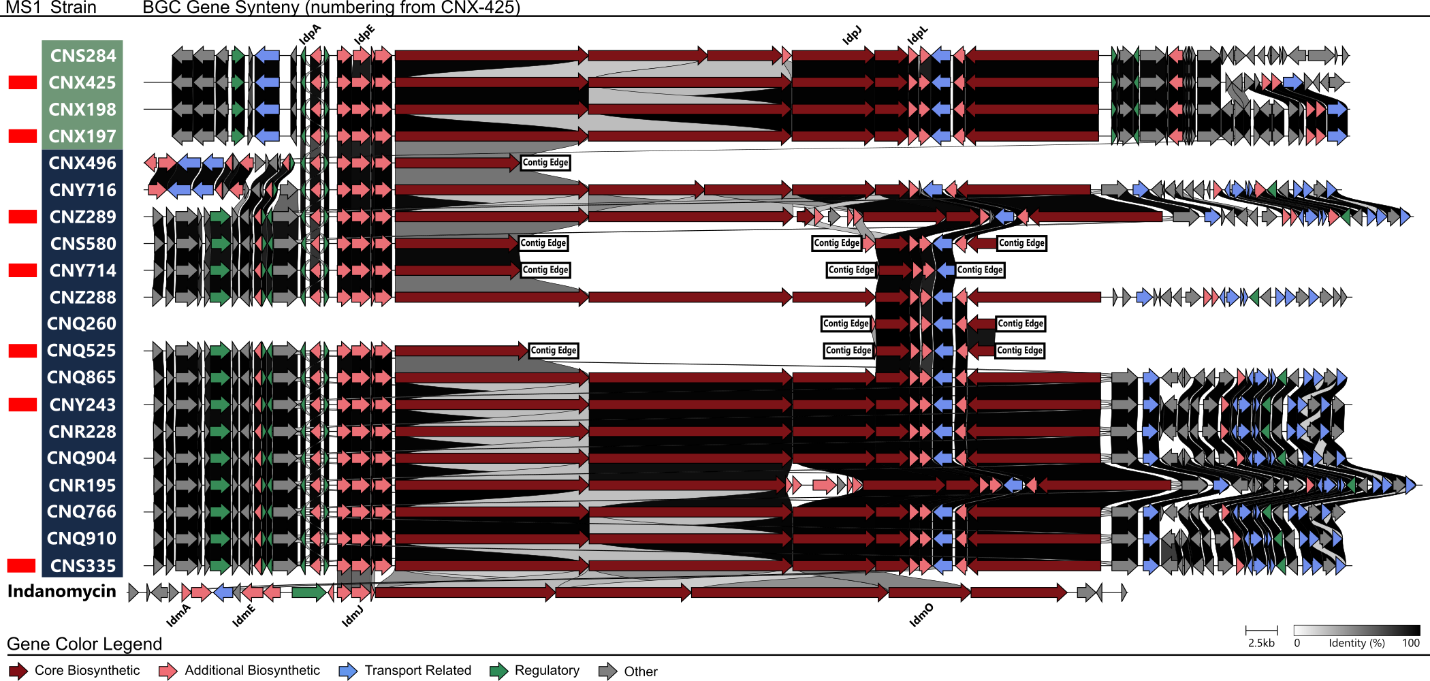

**Figure S6.** Synteny plots of 20 full and partial indanopyrrole BGCs found in MAR4 genomes. The indanomycin BGC is shown for reference. Red bars in the MS1 column indicate that indanopyrrole production was detected by MS in the small-scale survey. The strain column is colored (green or dark blue) to distinguish the two groups of strains that share >95% ANI.

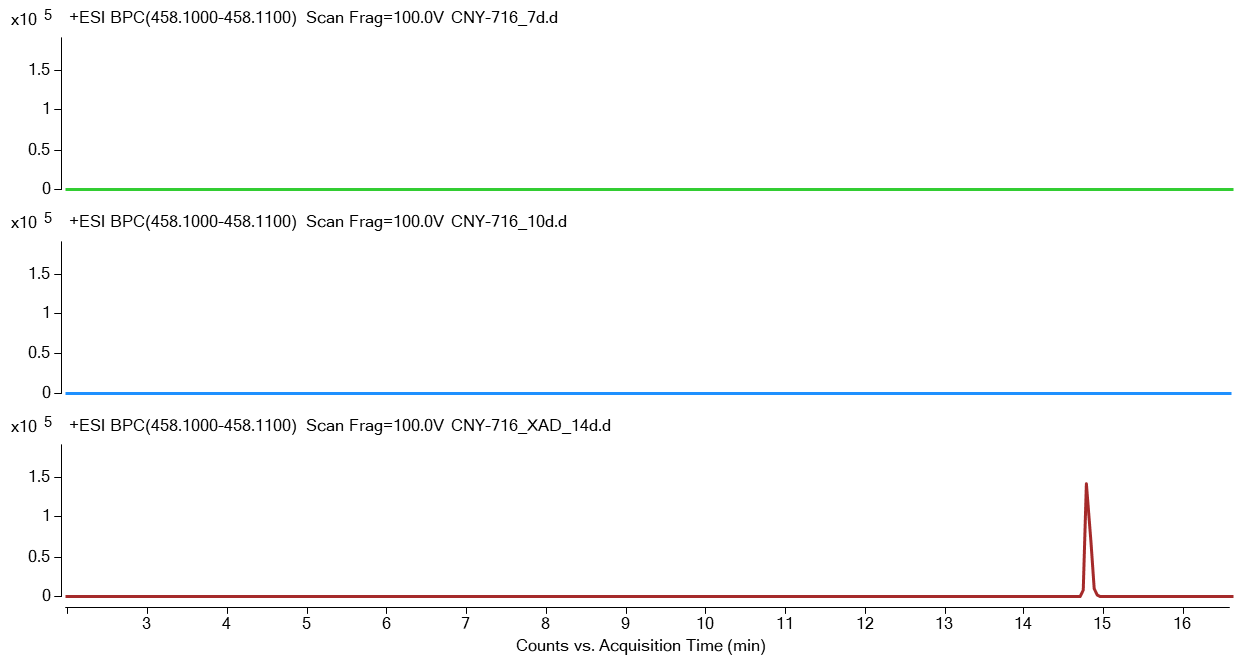

**Figure S7.** Production of indanopyrrole A (**1**) in *Streptomyces* sp. strain CNY-716. LCMS analysis of a 1 L culture over time. Extracted ion chromatograms (*m/z* 458.10-458.11) of EtOAc extracts after day 7 (top), day 10 (middle) and day 14 (lower) with production observed following the addition of XAD-7 resin on day 10.

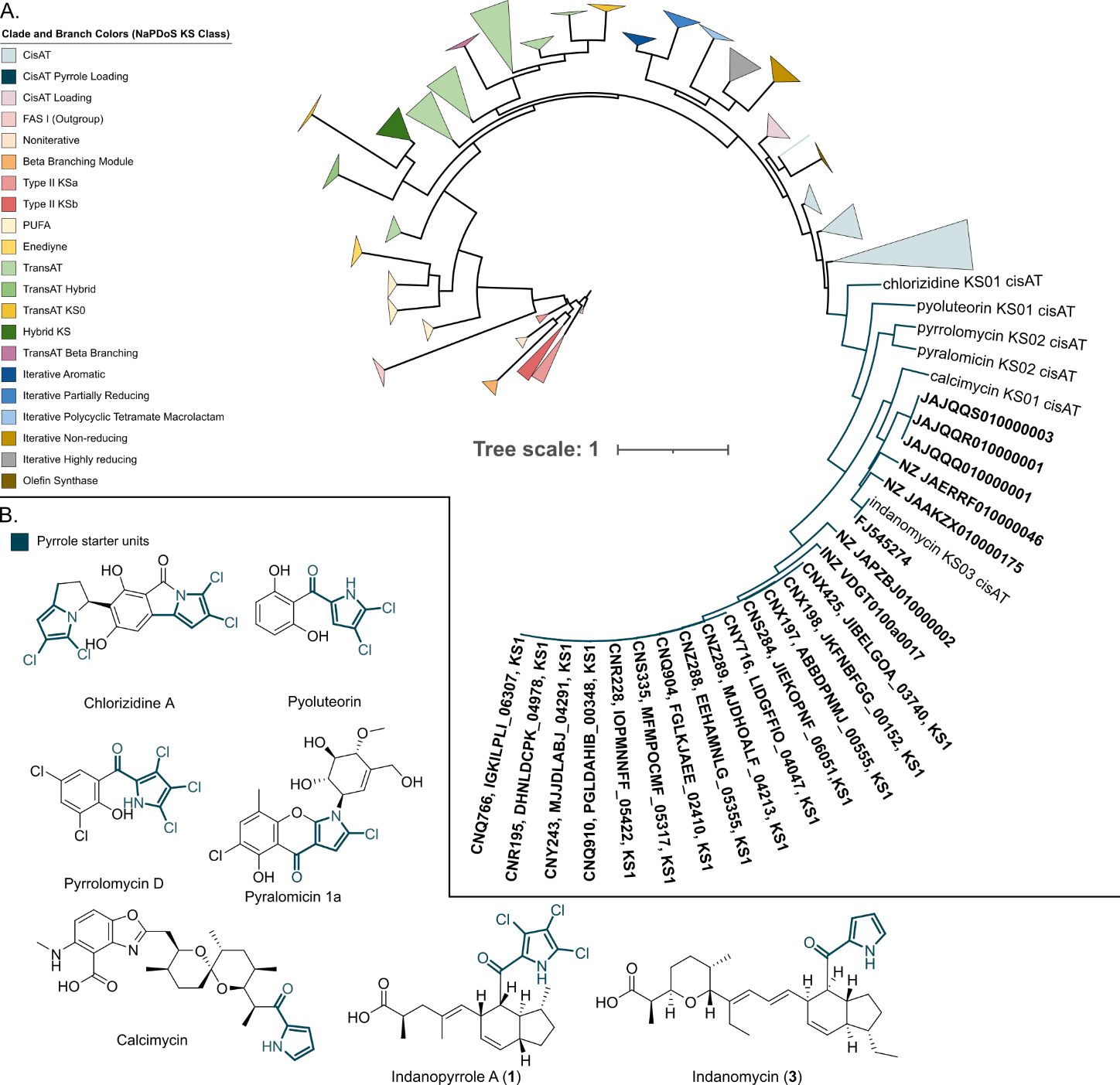

**Figure S8.** NaPDoS2 KS domain phylogeny. A. FastTree phylogeny from NaPDoS2 includes reference sequences from the NaPDoS2 database plus the KS domains from the loading modules of all idp and idm BGCs found (bold text). The idp and idm loading domains form a clade with those from BGCs that use pyrrole starter units (dark blue branches). B. Structures of compounds with pyrrole incorporating starter units, colored dark blue.

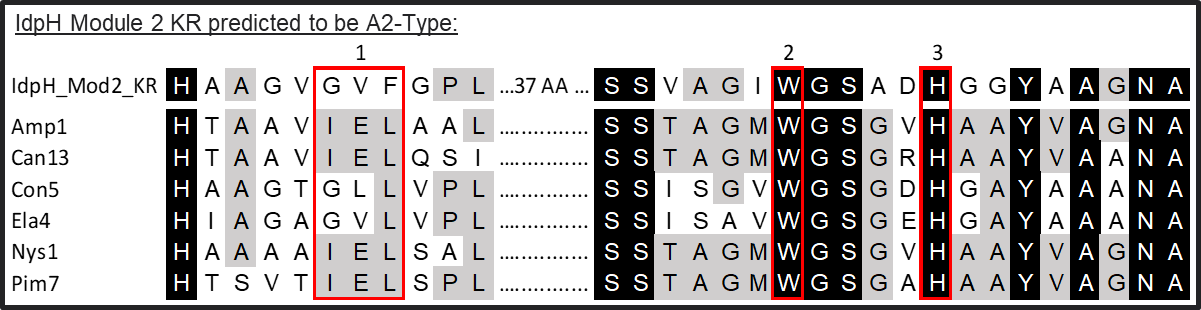

**Figure S9.** Ketoreductase sequence alignments. An alignment of the module 2 KR domain sequence from idpH and A2-type KR domains reveals the absence of the LDD motif (red box 1) associated with B-type domains and contains the active site tryptophan (red box 2) and histidine (red box 3) associated with A2-type domains, which are known to generate 2S-3S chemistry at the newly created stereocenter.

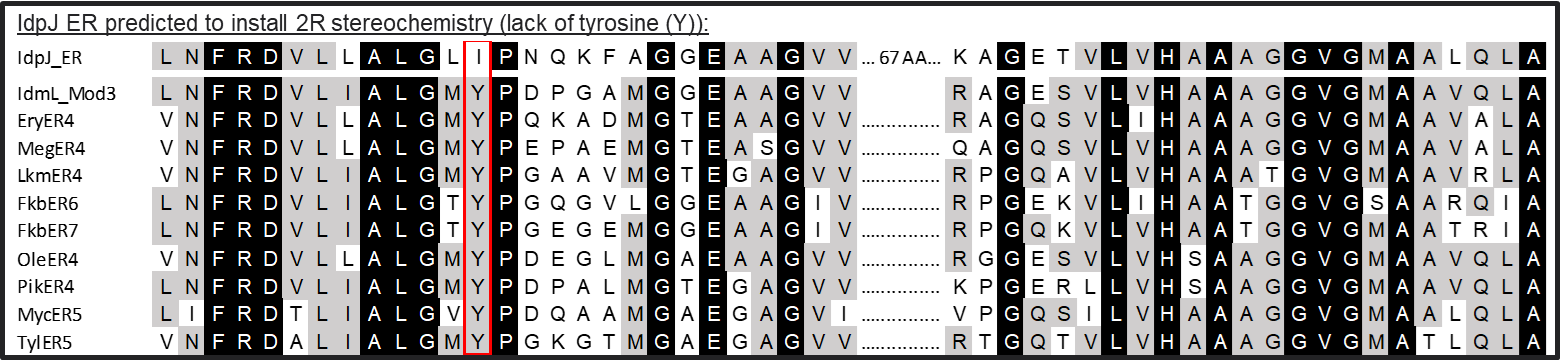

**Figure S10.** Enoylreductase sequence alignments. IdpJ is missing the active site tyrosine (red box) observed in ER domains that generate S stereochemistry, including that of idmL module 3.

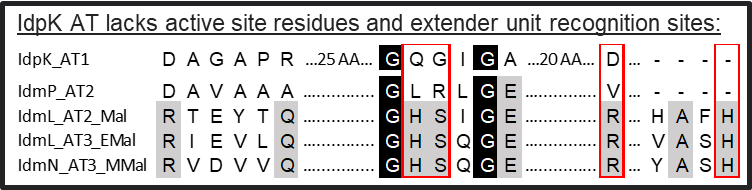

**Figure S11.** idpK acyltransferase (AT) domain alignment. The idpK_AT1 and idmP_AT2 AT domains are missing the active site residues involved in catalysis and extender unit recognition found in active AT domains included in the alignment (red boxes). IdpK_AT1 and idmP_AT2 are predicted to be inactive.

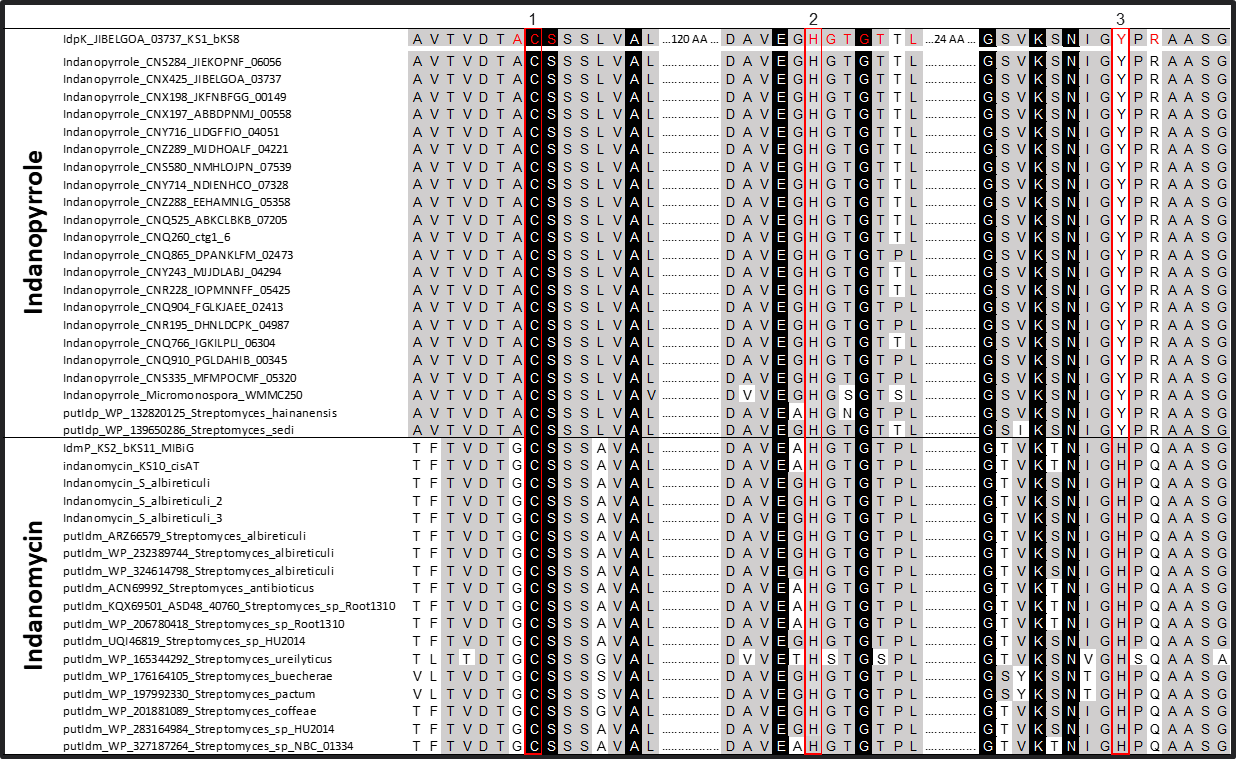

**Figure S12.** Terminal ketosynthase domain alignment from indanopyrrole producers and related BGCs. The active site histidine required for decarboxylation activity has been mutated to a tyrosine in all MAR4 indanopyrrole producers (red box 3). Residues in red font were shown to be within 5 Ångstroms of the docked indanopyrrole A molecule (Figure S14) and include the known catalytic residues.

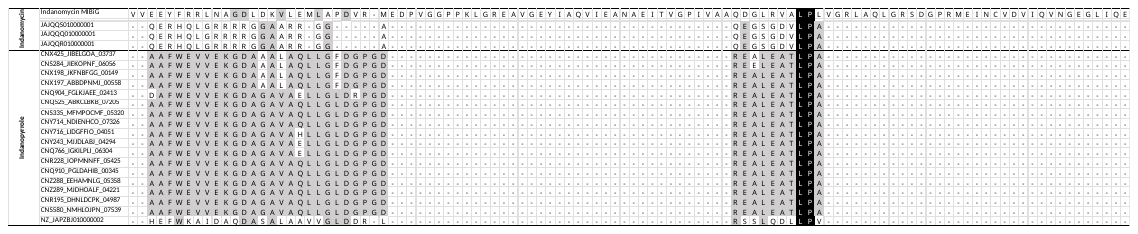

**Figure S13.** Indanomycin cyclase domain alignments. Only the Indanomycin BGC from MIBiG and DoBISCUIT databases contained the terminal cyclase domain, no other terminal PKS modules related to indanopyrrole or indanomycin found in this study contain this domain. Domain boundaries based on the indanomycin BGC on the DoBISCUIT database.^49^

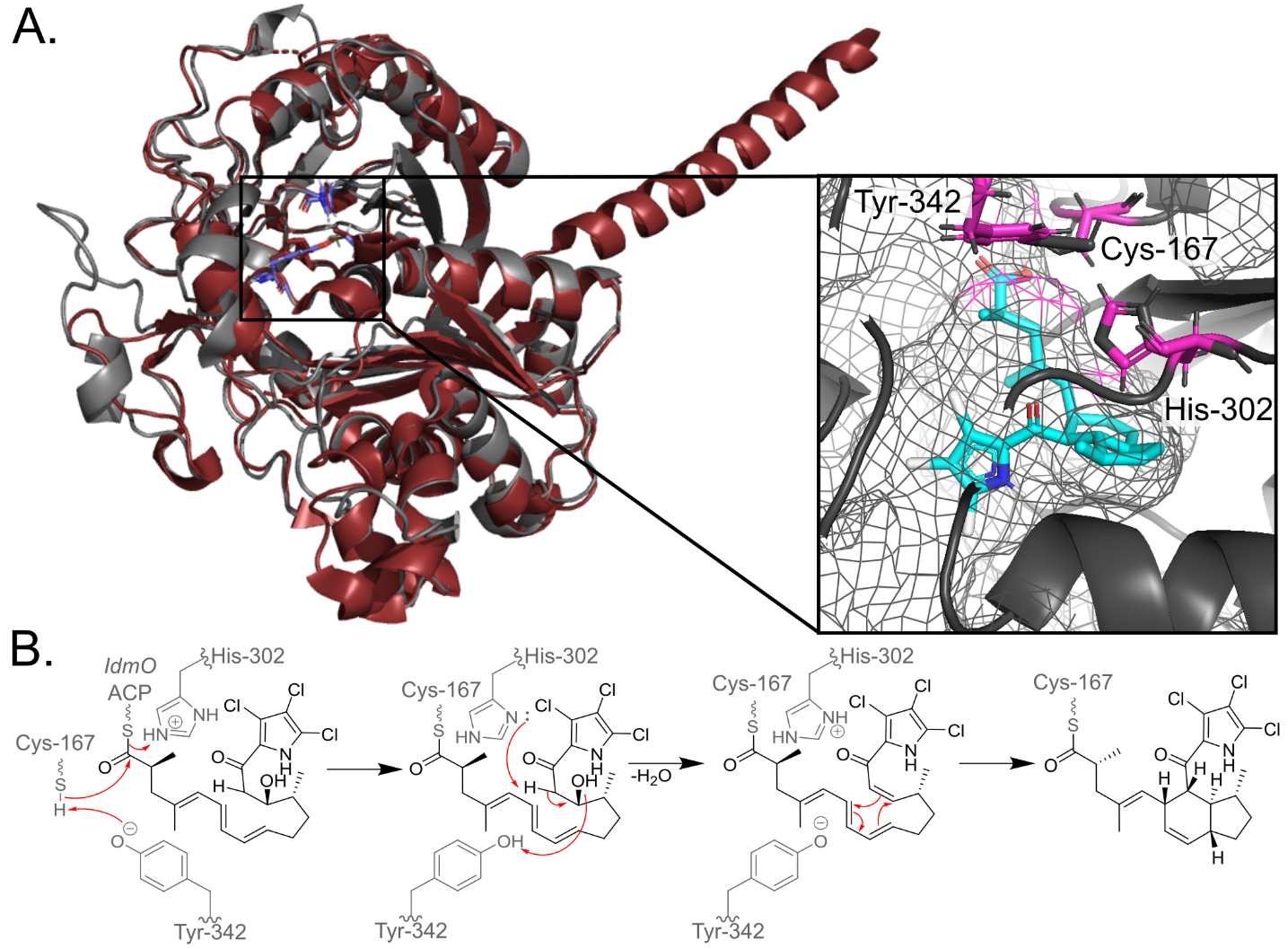

**Figure S14.** Modeling of IdpK against DEBS KS1. A. Colabfold model of IdpK KS (grey) matches well with the known 3D structure of DEBs KS1 (red). Inset: indanopyrrole A (cyan) modelled in the KS active site showing the alignment of cysteine-167 with the carboxylic acid moiety as expected in PKS biosynthesis. Histidine-302 is positioned above C14 in the indanopyrrole structure possibly allowing for hydrogen abstraction and a subsequent, spontaneous, Diels-Alder cyclization reaction to form the final product. Mesh surface indicates that the substrate binding pocket of IdpK is shaped in a way that could allow for a Diels-Alder reaction. B. Predicted biosynthetic mechanism for IdpK KS dehydratase activity and subsequent Diels-Alder reaction.

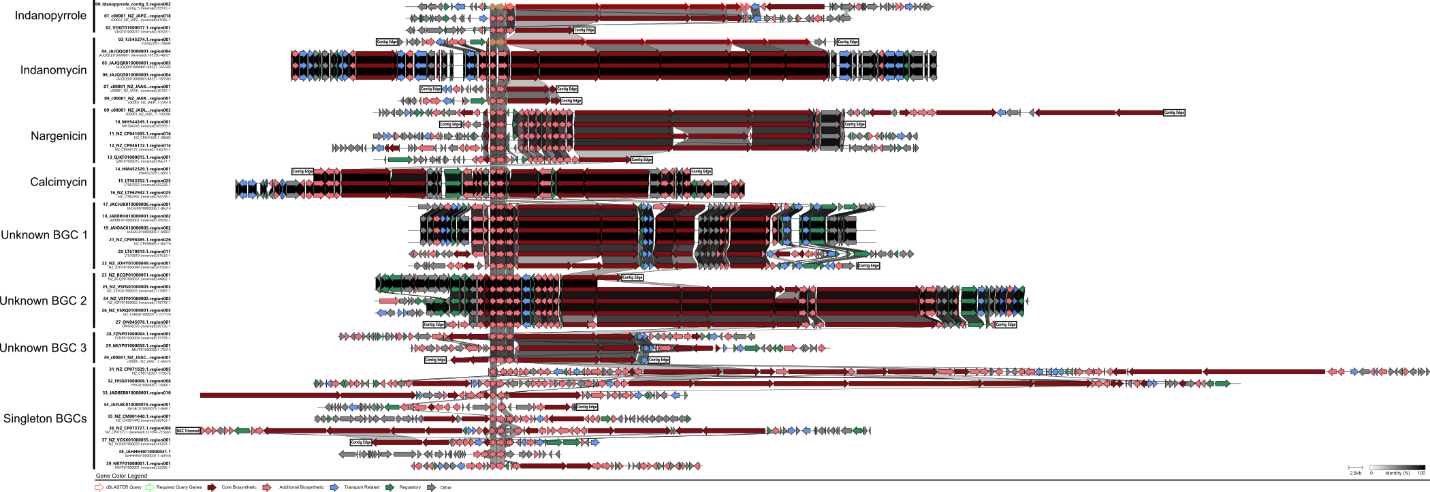

**Figure S15.** Idp and idm NCBI matches. cBlaster results filtered for the presence of at least one gene with >70% similarity to a PKS gene in idp or idm. The 40 BGCs were classified into seven BGC groups and nine singleton BGCs. BGC identities are listed on the right when known. Idp and idm are shown at the top.

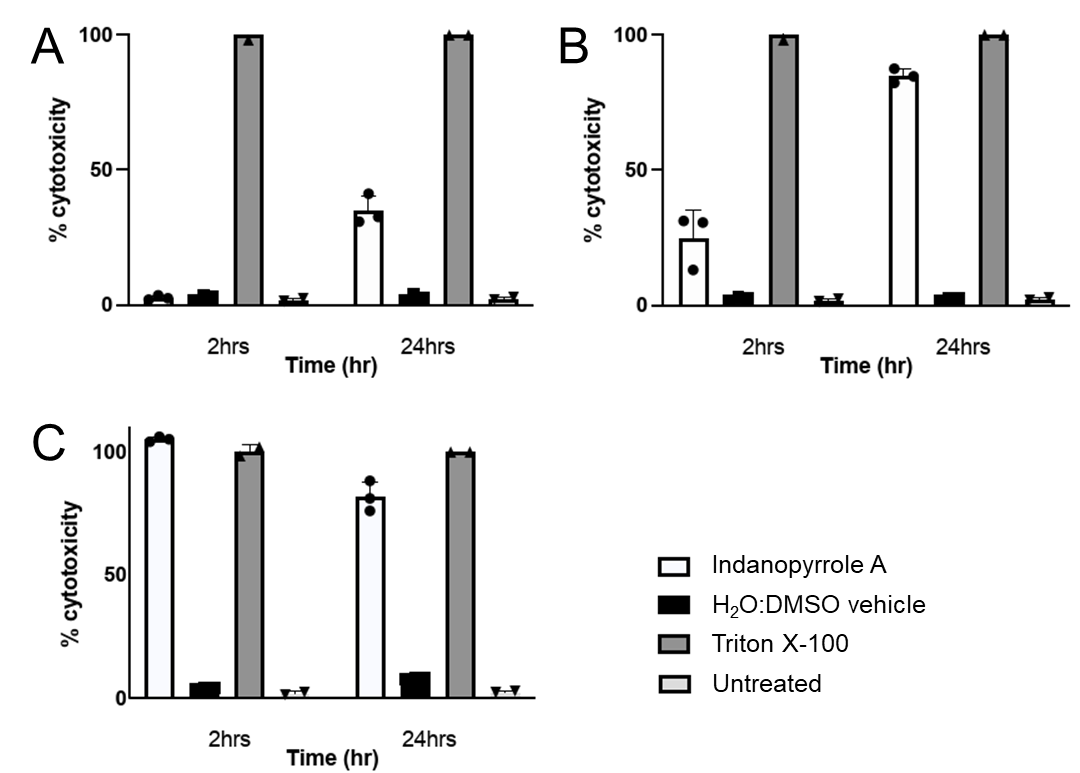

**Figure S16.** A549 cell line LDH cytotoxicity of indanopyrrole A (**1**). The compound was tested at 16, 32, and 128 µg /mL (panels A-C, respectively). Solvent vehicle (H_2_O:DMSO in 1:2 ratio) and untreated cells were used as negative controls. Cell lysis with Triton X-100 was used as a positive control. Cell toxicity was observed at 16 µg /mL after 24 hours.

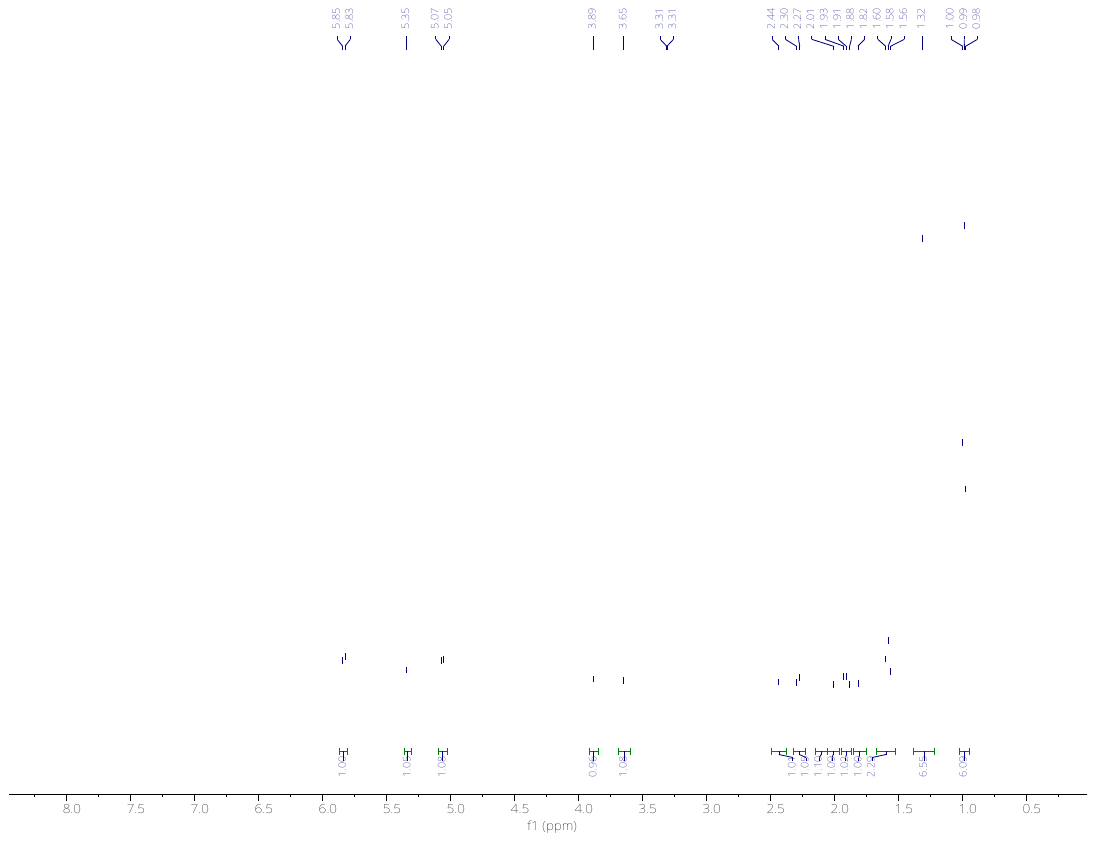

**Figure S17.** ^1^H NMR spectrum of indanopyrrole A (**1**) in MeOH-*d4*.

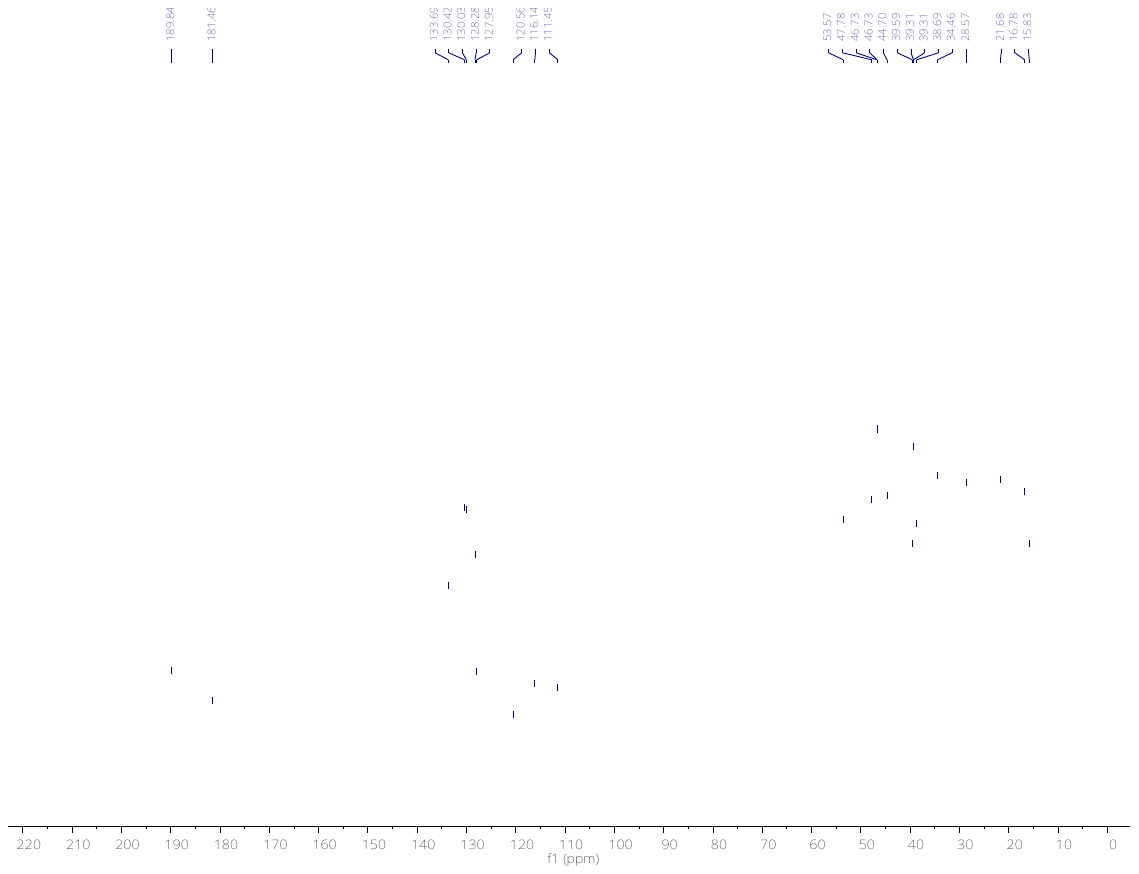

**Figure S18.** ^13^C NMR spectrum of indanopyrrole A (**1**) in MeOH-*d4* (Varian, 125 MHz).

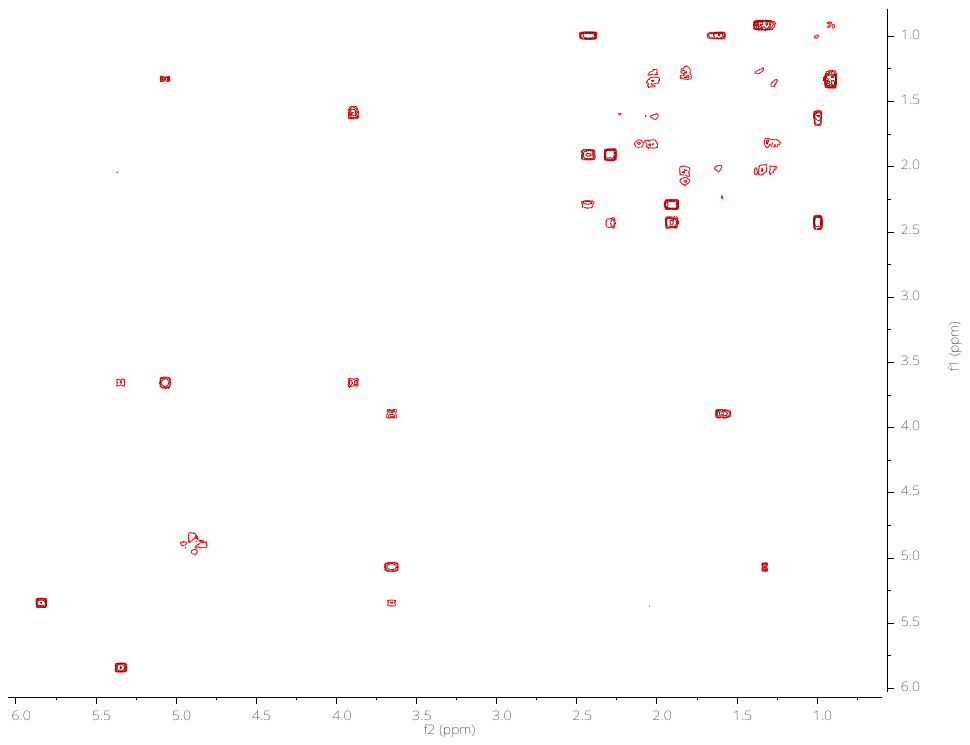

**Figure S19.** ^1^H-^1^H COSY spectrum of indanopyrrole A (**1**) in MeOH-*d4*.

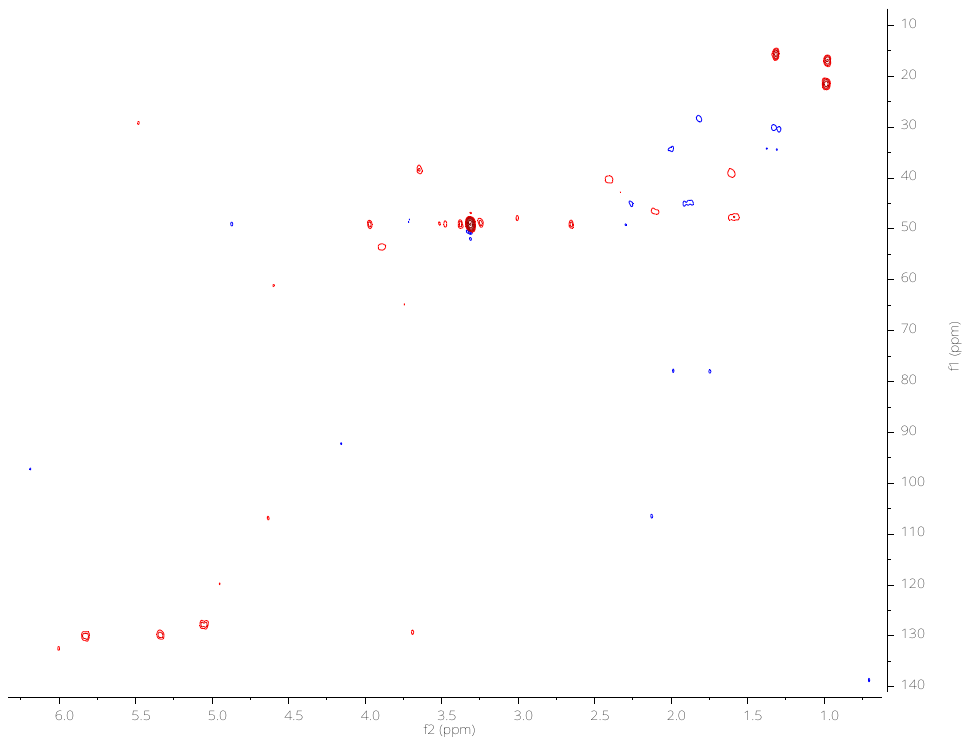

**Figure S20.** HSQC spectrum of indanopyrrole A (**1**) in MeOH-*d4*.

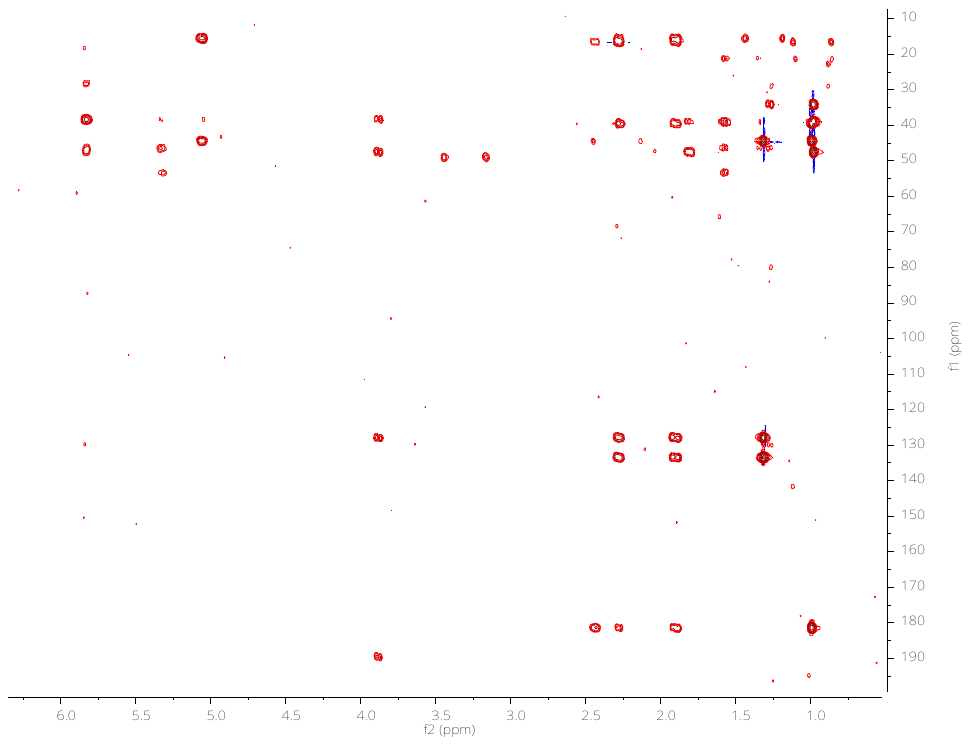

**Figure S21.** HMBC spectrum of indanopyrrole A (**1**) in MeOH-*d4*.

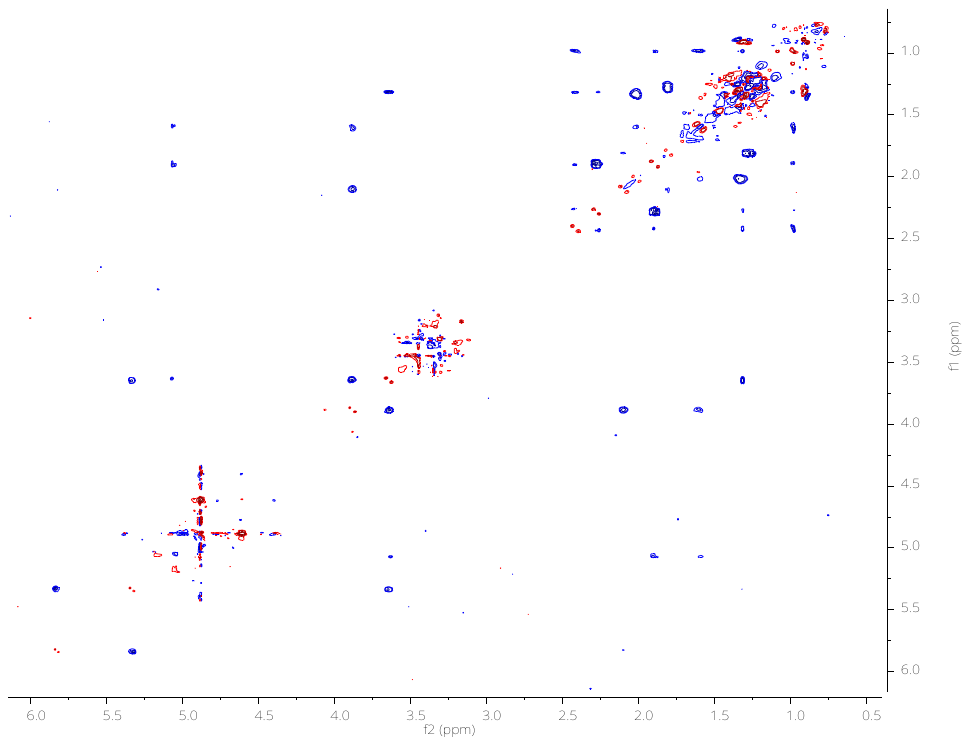

**Figure S22.** NOESY spectrum of indanopyrrole A (**1**) in MeOH-*d4*.

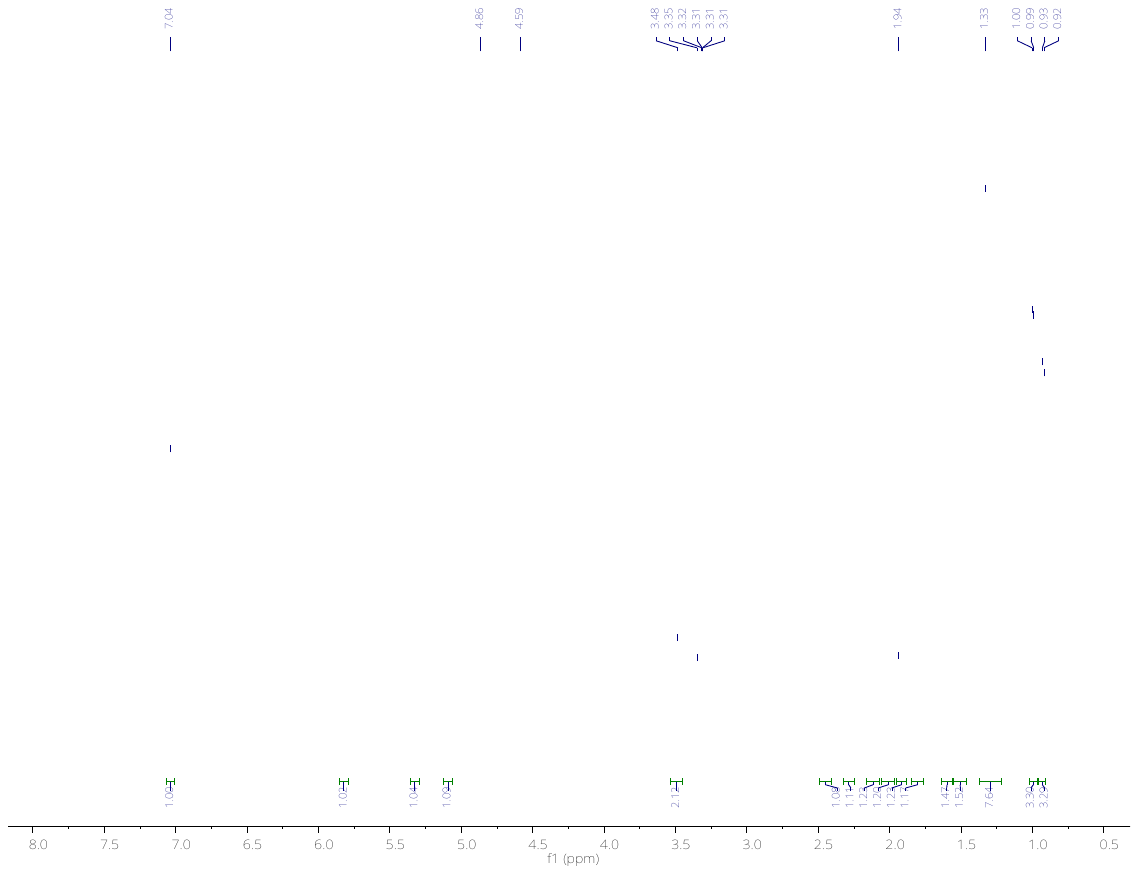

**Figure S23.** ^1^H NMR spectrum of indanopyrrole B (**2**) in MeOH-*d4*.

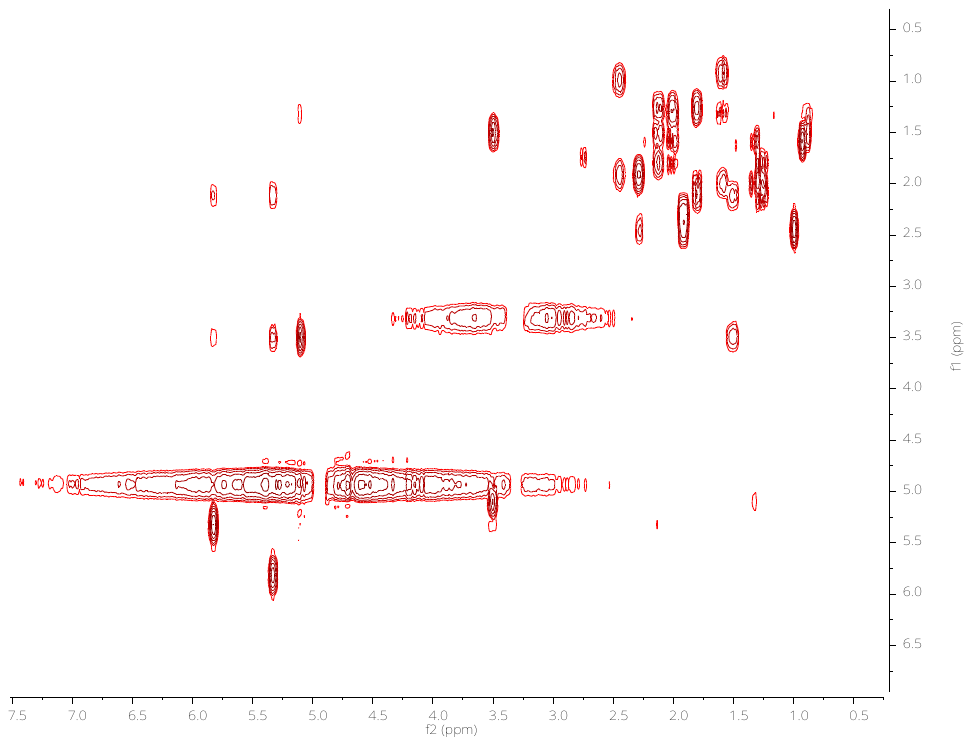

**Figure S24.** ^1^H-^1^H COSY spectrum of indanopyrrole B (**2**) in MeOH-*d4*.

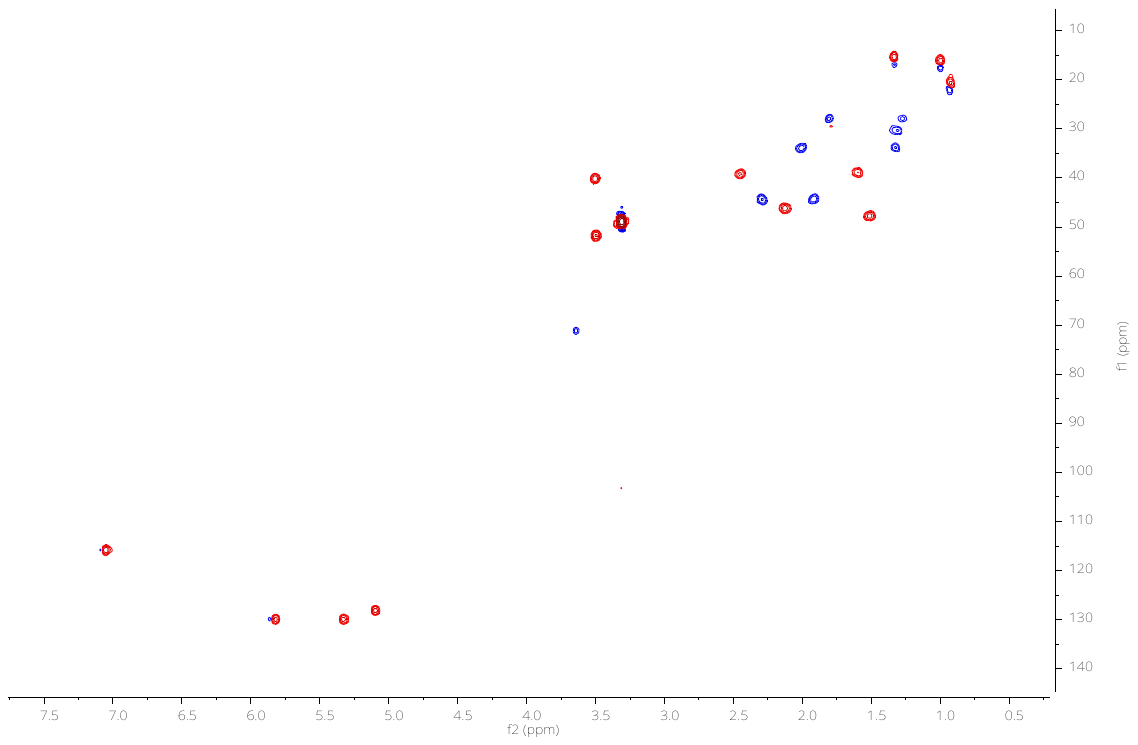

**Figure S25.** HSQC spectrum of indanopyrrole B (**2**) in MeOH-*d4*.

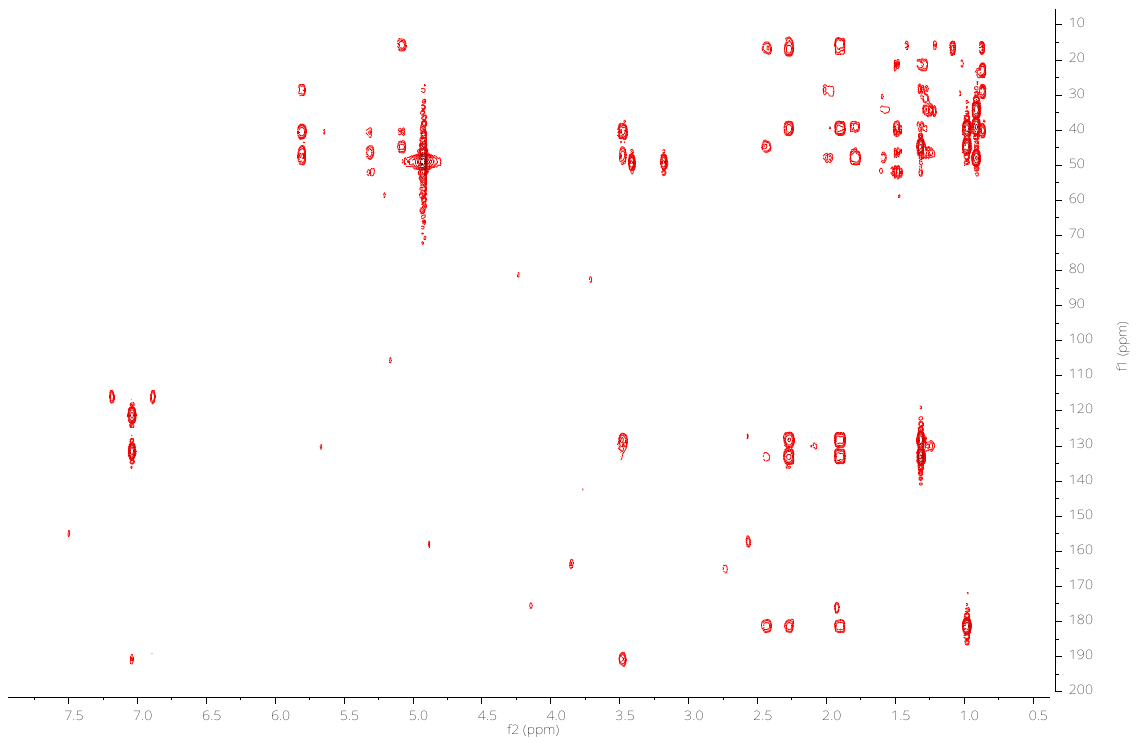

**Figure S26.** HMBC spectrum of indanopyrrole B (**2**) in MeOH-*d4*.

**Table S3.** Cartesian coordinates of **1** (minimal energy conformer).

0 1

C 1.24407833 1.78045416 2.07580351

C 1.54072911 0.31581622 2.31235657

H 2.32843990 0.24276790 3.06836467

C 2.36592991 2.72640079 1.97212673

O 0.11406945 2.19102463 1.84312991

N 2.13539864 3.91614811 1.32757735

H 1.25730142 4.13349025 0.84226881

C 3.27829468 4.61809648 1.22735499

C 4.29661427 3.90884963 1.84194565

C 3.71684100 2.71390301 2.31391776

Cl 4.58297545 1.50329024 3.18229683

Cl 5.93439907 4.39702601 1.99572698

Cl 3.34697504 6.13663152 0.44273459

C 2.15285814 -0.25327608 0.97085104

C 2.16584070 -1.77152434 0.94421638

H 2.70994189 -2.22022823 0.11551990

C 1.53969985 -2.54489520 1.82874508

H 1.55117812 -3.62697935 1.71183068

C 0.84868193 -1.95937093 3.01883632

H 1.58062871 -1.90700652 3.84392222

C 0.35336896 -0.53088713 2.76730681

C -0.38242509 -0.21822460 4.08512112

H 0.38376324 -0.01910080 4.84918110

C -1.07286552 -1.57499842 4.42325302

H -2.14276447 -1.51110691 4.19780408

H -0.99099441 -1.79377870 5.49240137

C -0.39392300 -2.66498337 3.55927697

H -1.03976378 -2.96109549 2.72426736

H -0.15477974 -3.57100863 4.12417482

C -1.38473060 0.93177414 4.08593412

H -0.90418454 1.90762826 4.00596855

H -2.08258202 0.83570821 3.24587573

H -1.97058987 0.91432854 5.01249018

H -0.39230692 -0.55582863 1.95814987

C 1.39885024 0.24003854 -0.24606046

H 0.36782327 -0.11062230 -0.30368635

C 1.82794017 1.03360597 -1.23651606

C 0.91954954 1.28127745 -2.42394831

H 1.46847775 1.10063722 -3.35720668

H 0.07537075 0.58551344 -2.40329555

C 0.36235825 2.72121105 -2.48479565

H 1.17237873 3.42474098 -2.69914376

C -0.71979563 2.85608598 -3.56273350

H -0.30439357 2.60128030 -4.54257116

H -1.55756819 2.18143176 -3.35995962

H -1.10255031 3.88003171 -3.61117743

C -0.18576005 3.12617445 -1.13622549

O -1.02704578 2.22888020 -0.62745930

H -1.10868988 2.40903768 0.32945208

O 0.10179343 4.16345700 -0.56681781

C 3.14185729 1.77070600 -1.26099366

H 3.65659528 1.60752095 -2.21616840

H 2.98763007 2.85301693 -1.16235781

H 3.81451678 1.47262028 -0.45435235

H 3.19456401 0.08430960 0.91528622

1 2 1.0 4 1.0 5 2.0

2 3 1.0 14 1.0 21 1.0

3

4 6 1.0 10 2.0

5

6 7 1.0 8 1.0

7

8 9 2.0 13 1.0

9 10 1.0 12 1.0

10 11 1.0

11

12

13

14 15 1.0 35 1.0 55 1.0

15 16 1.0 17 2.0

16

17 18 1.0 19 1.0

18

19 20 1.0 21 1.0 27 1.0

20

21 22 1.0 34 1.0

22 23 1.0 24 1.0 30 1.0

23

24 25 1.0 26 1.0 27 1.0

25

26

27 28 1.0 29 1.0

28

29

30 31 1.0 32 1.0 33 1.0

31

32

33

34

35 36 1.0 37 2.0

36

37 38 1.0 51 1.0

38 39 1.0 40 1.0 41 1.0

39

40

41 42 1.0 43 1.0 47 1.0

42

43 44 1.0 45 1.0 46 1.0

44

45

46

47 48 1.0 50 2.0

48 49 1.0

49

50

51 52 1.0 53 1.0 54 1.0

52

53

54

55
